# The Adolescent Functional Connectome is Dynamically Controlled by a Sparse Core of Cognitive and Topological Hubs

**DOI:** 10.1101/2025.03.14.643301

**Authors:** Jethro Lim, Ilias Mitrai, Prodromos Daoutidis, Catherine Stamoulis

## Abstract

Fundamental mechanisms that control the brain’s ability to dynamically respond to cognitive demands are poorly understood, especially during periods of accelerated neural and cognitive maturation, such as adolescence. Using a sparsity-promoting feedback control framework we investigated the controllability of the adolescence functional connectome. Critical feedback costs associated with a region’s control action on itself and the rest of the brain were estimated using resting-state fMRI data from an early longitudinal sample in the Adolescent Brain Cognitive Development (ABCD) study (n = 1394; median (IQR) age = 10.1 (1.1) years at baseline and 12.1 (1.1) years at follow-up). A highly reproducible, core set of predominantly highly connected regions retained their control action over the connectome under high feedback costs. They included posterior visual areas, retrosplenial cortex, cuneus and precuneus, superior parietal lobule, temporal ventral cortex and dorsolateral and lateral prefrontal cortices, i.e., both developed and developing brain regions. These regions were central to the topological organization of the connectome, consistently engaged during spontaneous coordination of resting-state networks, and overlapped with cognitive and topological brain hubs that play ubiquitous roles in cognitive function and the organization of the connectome. Also, most received (integrated) and distributed approximately equal amounts of neural information. These regions’ control action was developmentally stable, i.e., critical feedback costs did not change significantly during puberty, suggesting that, despite ongoing maturation and topological changes in the adolescent brain, fundamental mechanisms of system controllability may be well developed to facilitate information processing and response to cognitive demands.

## 1. INTRODUCTION

The brain’s ability to respond to myriads of external inputs and complex cognitive demands critically depends on its capacity to transition between states (physiological, cognitive, topological and/or dynamic) that are accompanied by specific patterns of interaction between its regions (Myer-Lyndenberg et al, 2002; Deco et al, 2011; Cocchi et al., 2013; Tognoli and Kelso, 2014; Breakspear, 2017; Vidaurre et al, 2017; Shine and Poldrack, 2018; Taghia et al, 2018; Kringelbach and Deco, 2020). These transitions occur at multiple time scales that depend on processing demands and type of state (for example long scales associated with arousal, wakefulness, or sleep, i.e., (order of minutes or hours), versus short scales associated with dynamic states (order of milliseconds)). Mechanisms that facilitate brain state transitions, and the dynamic characteristics of individual states (such as their stability), are incompletely understood (Hansen et al., 2015).

Various methods and models have been used to estimate human brain dynamics at temporal scales resolved by individual neuroimaging modalities (for example millisecond resolution of EEG vs (sub)second resolution of fMRI), and elucidate their encoding of brain states (Kringelbach and Deco, 2020). Prior work has shown that even at rest, intrinsic dynamics are characterized by discrete states of synchrony (dynamic connectivity) that may be associated with cognitive performance and introspective processes that may be associated with introspective and metacognitive processes (Marusak et al, 2017, 2018; Di et al, 2023; Ye et al, 2024; Fu et al, 2024).

Recently, the field has used tools from control theory to elucidate the brain’s controllability, and thus mechanisms that underlie its ability to switch between dynamic states. Some of these tools have been applied to graph (network) representations of brain circuits, with the goal to investigate the brain’s response to control inputs in specific regions (Pasqualetti et al., 2014; Gu et al., 2015; Betzel et al., 2016; Lee et al., 2019; Parkes et al., 2023). The majority of prior brain studies using control theoretic tools have used an open-loop framework and have shown that state transitions may be controlled by a small set of brain regions that are active in a particular state (Gu et al., 2015, 2017). In addition, highly connected regions form a ‘core’ network (van den Heuvel & Sporns, 2011), that supports the minimization of control energy required for the brain to transition between states (Betzel et al., 2016) and optimize cognitive function (Cui et al., 2020). Within this framework, control energy has also been correlated with temporal changes in functional connectivity (Deng et al., 2022). Recent translational work has also used the open-loop control framework to elucidate changes in brain dynamics and controllability associated with neuropsychiatric disorders, such as schizophrenia and psychosis, where networks lose their cost efficiency as the brain remains in energy-demanding states (Braun et al., 2021, Zoller et al., 2021, Parkes et al., 2021). Network controllability has also been examined as a predictor of aphasia recovery after stroke (Wilmskoetter et al., 2021) and psychosis spectrum symptoms (Parkes et al., 2021).

In contrast to the open-loop framework, a closed-loop control framework assumes that internal processes control the system’s dynamics, and feedback information drives adjustments to the controller’s action, in order to optimize the system’s function. This type of control framework has been used in brain stimulation, brain-machine interface and neurofeedback (Wright et al., 2016, Zrennar et al., 2016, Sitaram et al., 2017, Skarpaas et al., 2019, Zhang et al., 2023). These applications specifically seek to optimize either the brain’s dynamic response (e.g., to stimulation) or motor behavior through feedback signals. Few, if any, studies have used this framework to elucidate how neural dynamics are topologically controlled, through latent processes, especially during development when the brain undergoes profound reorganization, and its circuitry is incompletely maturated and suboptimally wired.

The developed brain has a small-world and rich-club topological organization that facilitates efficient local (domain-specific) computation and global (domain-general) integration through sparse long-range connections (van den Heuvel and Sporns, 2011, Bassett and Bullmore, 2017). Its modular topology is thus characterized by sparsity (Eavani et al., 2015), and a sparsity-promoting, feedback control framework is conceptually appropriate for studying the controllability of brain networks (Lin et al., 2013, Constantino et al., 2019). In such a framework, the controller’s action is spatially sparse, and is optimized through balance between performance and feedback costs, a process that is likely well-aligned with the action of biological mechanisms underlying the controllability of brain networks. In recent work, we have applied this control framework to investigate the controllability of macaque structural networks (Mitrai, et al. 2021) and in preliminary studies of human functional networks in adolescence (Mitrai et al., 2023, Lim et al., 2024).

In this study, we investigated the controllability of maturating brain networks in adolescence. We hypothesized that despite its suboptimal and evolving organization, the developing connectome is controlled by a relatively sparse set of regions that play a critical role in controlling its dynamics. To test this hypothesis, we applied a sparsity-promoting feedback control framework to intrinsic networks from a sample of ∼1,400 adolescents from the Adolescent Brain Cognitive Development (ABCD) study (Casey et al, 2018) with early longitudinal neuroimaging data (baseline (ages ∼9-10 y) and two-year follow-up (ages ∼11-12 y), spanning pre/early to late puberty. We investigated associations between estimated feedback control costs and regional connectome characteristics, including connectedness, information flow, and dynamics. Finally, we assessed relationships between region-specific feedback control costs and demographic and other participant characteristics, especially pubertal stage, in order to elucidate the impact of brain development on the connectome’s internal control.

## 2. METHODS

### 2.1 Sparsity Promoting Feedback Framework

The control framework used in the study is described in detail in prior work (Lin et al., 2012; Mitrai et al., 2023). Briefly, the brain is represented as a linear time-invariant system described by:

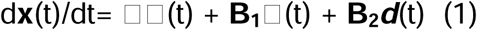

where is the system state (generally corresponding to a region), ***u*** is the control input (activated internally rather than externally), ***d*** are the external input(s), describes the internal dynamic behavior of the system, and **B_1_** and **B_2_** are matrices that map the control and external inputs to the system state. The control input is related to the state by:

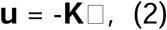

where **K** is the feedback gain matrix. Each element (**i,j**) of **K** represents the feedback control action that region **j** exerts on region **i**. **K** is the target of optimization and is obtained by penalizing its non-zero entries while maximizing control performance, resulting in the following optimization (controller synthesis) problem:

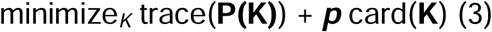

where ***p*** captures the feedback cost, card(.) is cardinality, and **P(K)** is related to **A** and **K** by:

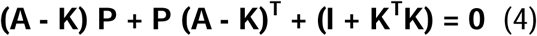

As *p* increases, the cardinality of K decreases and thus its sparsity increases.

This optimization problem was solved using the alternating direction method of multipliers (ADMM) algorithm (Boyd et al., 2011; Lin et al., 2012), restricting the range of ***p*** values to *^p^*^lJ [10^*^−^* ^4, 30]^. The upper bound was selected empirically, by examining the number of regions retaining their control action as *p* increases, i.e., the cardinality of **K** (see Figure S5 for an example). At *p* > ∼20, the cardinality of K remained almost constant. Within the selected range, two critical values were of specific interest, *p* at which a region’s control action does not depend on any other region (i.e., the feedback gain matrix becomes diagonal), and a critical transition *p* value at which a region no longer exerts its control action over the network (or retains its action if p has reached the upper limit).

### 2.1 Participants

Resting-state (rs) fMRI data from 1394 youth (53.7% female, 53.7% white non-Hispanic, 21.7% Hispanic) in the ABCD study were analyzed. Median age (interquartile range (IQR)) was 10.1 (1.1) years at baseline and 12.1 (1.1) years at follow-up. All participants had early longitudinal data (i.e., two assessments, at baseline and 2-year follow-up). The sample was selected based on a requirement of two high-quality rs-fMRI runs (each 5 min long) at both assessments for replication analyses. Data quality in a particular run was assessed based on median connectivity of the entire connectome (which is inherently low at rest) and the number of frames censored for motion (based on a displacement threshold of 0.3 mm). In the selected runs, median percent of frames censored for motion was 0.8%-1.3% at baseline, and 0.5%-0.8% at follow-up.

### 2.2 fMRI Data Processing

All analyzed rs-fMRI data (from the ABCD release 4.0) had been minimally preprocessed by the Data Analysis, Informatics & Resource Center (DAIRC; Hagler et al, 2019), and were further processed using the Next-Generation Neural Data Analysis (NGNDA) pipeline. Processing details are provided in Brooks et al, 2021. Briefly, each participant’s fMRI data were corrected for motion, coregistered to their T1 structural MRI, normalized to NMI space, denoised to suppress artifacts, and harmonized across scanners (fMRI had been acquired at 21 sites of the ABCD, using 3.0T Siemens, GE and Philips scanners). Repetition time (TR) was 0.8 s across scanners. Voxel-level data were parcellated using 3 atlases (cortical, subcortical and cerebellar), resulting in 1088 parcels (1000 cortical, 54 subcortical and 34 cerebellar; Schaefer et al., 2018, Tian, 2020, Diedrichsen, 2009). To facilitate computationally tractable control analyses, the higher resolution data (1088 parcels) were then further downsampled to 100 regions based on anatomical boundaries and the delineation of large resting-state networks identified in Yeo et al., 2011. In secondary analyses that assessed the impact of spatial resolution on the results, a second parcellation using 300 cortical parcels (also based on Schaefer et al, 2018) was used, resulting in a total of 388 regions.

Connectivity matrices were estimated using peak pairwise signal cross-correlation (in the range 0-1) as a measure of connection strength. Each matrix was thresholded using assessment-specific, cohort-wide statistical thresholds (the same threshold was used for both runs within each assessment). Given that the brain is overall weakly correlated at rest, with a relatively low number of active functional connections, a threshold equal to median + 1.5*IQR (moderate outlier threshold) was selected. Values below the given threshold were set equal to 0, resulting in relatively sparse adjacency matrices. Secondary analyses were also conducted using a less conservative threshold, corresponding to the cohort-wide 75th percentile of cross-correlation values (which was also approximately equal to brain-specific proportional thresholds, which retained 25% of connections, and a percolation-based threshold estimated independently (Brooks et al, 2021)).

Before solving the control problem, each adjacency matrix **A** was normalized using its maximum eigenvalue λ_max_, to ensure that at time t = 0 it represented a stable dynamic system:

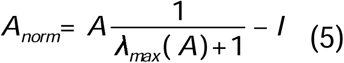

### 2.3. Regional topological characteristics

#### a. Net information flow

Flow of information through individual regions was estimated from fMRI time series using a variant of the Granger causality method (Chen et al, 2019). The optimal lag Granger-Geweke test (Geweke, 1982) was used to measure information flow between pairs of time series, based on their optimal lag, obtained by minimizing the Akaike Information Criterion (AIC) value. Two test statistics were further considered, the F_i->j_ value, representing a directed edge from region i to region j, and the corresponding p-value, used to assess the significance of the directionality. Only significant directed regions were considered. Net flow was calculated as the difference between information in and out of a region.

#### b. Regional connectedness (node degree) and dynamic engagement

Degree was estimated from the adjacency matrix as the total number of a region’s connections (edges). In addition, time-dependent adjacency matrices (estimated by thresholding corresponding correlation matrices, obtained using a 11-point sliding window (∼9 s) with a one-point increment) were used to estimate each region’s time-varying degree (Lim, et al, unpublished). Time points corresponding to frames censored for motion were then excluded, and each region’s degree was normalized by the brain-wide maximum estimated degree (across regions and time points). If a region’s (normalized) degree was greater or equal to a threshold of 0.5, the region was considered *engaged*. Frequency of region engagement was defined as the number of time points in which a region’s degree met or exceeded this threshold.

### 2.4 Statistical Analysis

Linear mixed effects regression models were developed to investigate associations between estimated feedback control costs and pubertal stage. Models included adjustments for sex, race/ethnicity (dichotomized as white non-Hispanic vs racial/ethnic minorities, given small samples and thus limited statistical power of individual racial categories), family income and BMI z-score (stratified by sex). These parameters were also independently examined in models to identify links between demographic and other participant characteristics and estimated costs. Models combined data from both baseline and two-year follow-up, and a random intercept and slope were included in the models to account for each participant’s repeated measurements. All statistical analyses accounted for sampling differences across the 21 ABCD sites, using propensity weights provided by the ABCD. Models were also adjusted for time of fMRI scan (rounded to the nearest hour, and thus represented by a 0-23 variable), to account for differences in topological parameters resulting from differences in the time of day at which participants were scanned (Hu et al, 2023), and percent of frames censored for motion (Brooks et al, 2021). All analyses assumed a significance level α = 0.05, and p-values were corrected for False Discovery Rate (FDR; Benjamin and Hochberg, 1995).

### 2.5 Similarity of spatial cost distributions

To compare the spatial distributions of control costs across resolutions, between runs, and between thresholds, two similarity measures were estimated, cosine similarity and the Barroni-Urbani (BU) coefficient. No resampling was necessary when comparisons were made at the same resolution. However, across the two resolutions of interest (100 vs 388 regions), estimated costs were mapped back to the original 1088 regions. Then, pairs of vectors (each with 1088 elements) were compared, using cosine similarity. In addition, to specifically compare the spatial patterns of regions associated with high control costs at the two resolutions, brain- and resolution-specific thresholds were imposed on weighted vectors, to binarize them and compare them using the BU coefficient. Two thresholds were selected, corresponding to the 75th and the 90th percentile of control costs at which regions no longer maintain their control action over the network.

## RESULTS

### 3.1 Distribution of feedback control costs across participants and brain regions

The distribution of costs at which individual regions maintained their control action over the network (or lost their action if costs are lower than the upper limit of the examined range) are shown in Figure 1, for baseline (a) and 2-year follow up (b). These costs were estimated from the best-quality fMRI run, and adjacency matrices thresholded based on the moderate outlying peak cross-correlation value. Corresponding findings for feedback costs at which regions become self-controlled (i.e., their dynamics do not depend on the control action of other regions) are shown in supplemental Figure S1. The spatial distribution of median (across participants) costs associated with self-control and loss of control action, respectively, is shown in Figure 2. Within the range of feedback costs associated with self-control ([min, max] = [<0.01, 0.30]), higher values were estimated in bilateral posterior areas overlapping with the visual peripheral network, bilateral superior parietal lobules (SPL), and bilateral dorsolateral prefrontal areas that partly overlapped with the salience, but more extensively with the frontoparietal control and default-mode (DM) networks. Within the range of feedback costs associated with regions maintaining (or losing) their control action over the network (median over participants of [min, max] = [8.58, 30.0]), the highest critical feedback costs were estimated in left dorsolateral prefrontal cortical areas, overlapping with the frontoparietal control network. In addition, high feedback costs were estimated in other bilateral frontal regions, overlapping with the DM and salience networks, as well as in posterior visual peripheral areas, bilateral SPL, the left ventral temporal area (part of the inferior temporal gyrus), overlapping with the frontoparietal control network, the right precuneus and right retrosplenial cortex. Overall, most of these regions had high critical feedback costs in both regimes of interest, and belonged to primarily 3 resting-state networks, the DMN, frontoparietal control, and visual peripheral network, i.e., a combination of developing (DMN and control) and developed (visual peripheral) networks.

**Figure 1:**
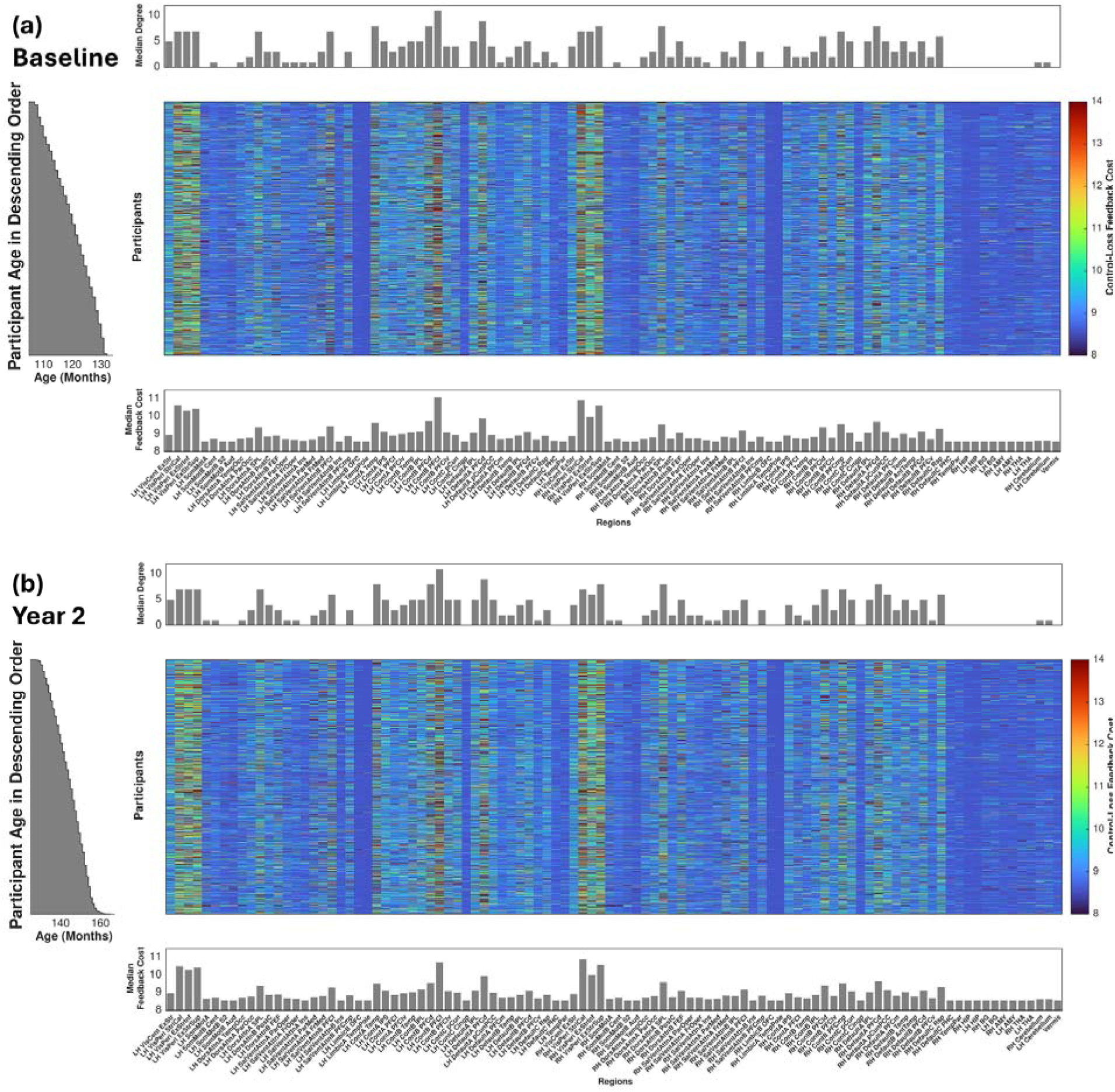
Feedback costs at which regions lose their control action on the network, as a function of participants (y-axis) and analyzed regions (x-axis). The color range corresponds to cost values, estimated from the best-quality run at baseline (a) and two-year follow up (b). Participants are sorted by age. Median (over participants) node degree for each analyzed region is shown in the bar plot above each heat map, and median feedback costs in the bar plot below each heat map. All parameters were estimated using adjacency matrices obtained by thresholding corresponding connectivity matrices, based on the moderate outlying peak cross-correlation value.

**Figure 2.**
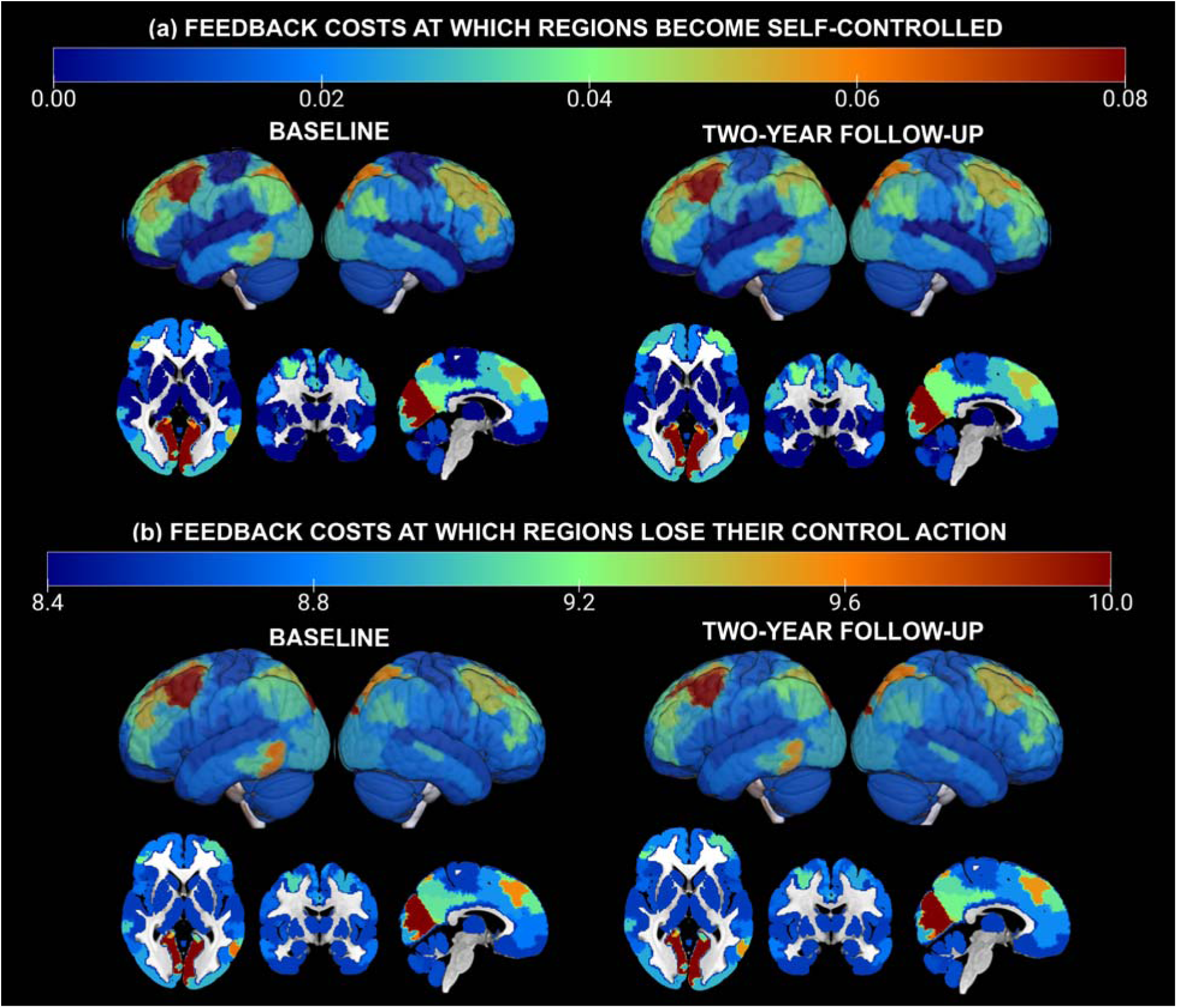
Spatial distribution of median (across participants) feedback costs at which (a) a region’s control action depends only on its own dynamic state, and (b) a region loses its control action over the network. Distributions are shown separately for baseline (left) and two-year follow up (right), and two-dimensional (horizontal, coronal, sagittal) slices and three-dimensional volumes are superimposed. Colors correspond to feedback cost values.

#### 3.1.1 Effects of connectivity matrix thresholding on control cost estimates

Secondary analyses examined critical feedback costs estimated from adjacency matrices that had been obtained with a less conservative threshold (the 75th percentile of cross-correlation values). Results are summarized in Figures S2-S4 (heat maps for costs as a function of participant (S2) and region (S3), and spatial distribution of these costs (S4)). There was substantial spatial overlap of high critical control costs with those estimated using a more conservative threshold, but cost values were overall higher with the less conservative threshold. Median cosine similarity between critical costs estimated based on the two thresholds was high across assessments (>0.87). Similarly, at both assessments, the spatial distributions of high control costs (top 25% and top 10%) was also consistently high (BU ≥0.84). These findings suggest overall invariance of the results to thresholding.

#### 3.1.2 Effects of fMRI run on control costs estimates

To assess the robustness of the results, the analyses were repeated using the second best-quality run, and estimated control costs were compared. At both assessments, cosine similarity of self-control costs estimated from the two runs was moderate (median (IQR): 0.65 (0.25) - 0.66 (0.25), and similarly for critical costs at which regions lost their control action (median (IQR): 0.48 (0.35) - 0.50 (0.34)). Similar similarity was estimated when top 25% and 10% of critical costs were compared between runs in each assessment (median (IQR) BU: 0.57 (0.24) - 0.63 (0.19)). These statistics reflect moderate similarity of spatial control cost distributions estimated from distinct rs-fMRI runs.

#### 3.1.3 Effects of spatial resolution on control cost estimates

To assess the impact of the spatial resolution at which the control costs were estimated, the analyses were repeated using a higher resolution parcellation (388 regions). Only the best-quality run was analyzed at this resolution. The spatial distribution of high critical costs and corresponding distribution of region degree at the two resolutions as shown in Figure 3. Cosine similarity between critical cost estimates was moderate (median (IQR): 0.45 (0.18) at baseline, and 0.44 (0.19) at follow up). Similarity of spatial patterns of highest 25% and 10% costs was also moderate at both assessments (median (IQR) BU: 0.63 (0.15) - 0.67 (0.10)). These statistics suggest at least partial correlation of control cost estimates with the spatial resolution of their assessment.

**Figure 3:**
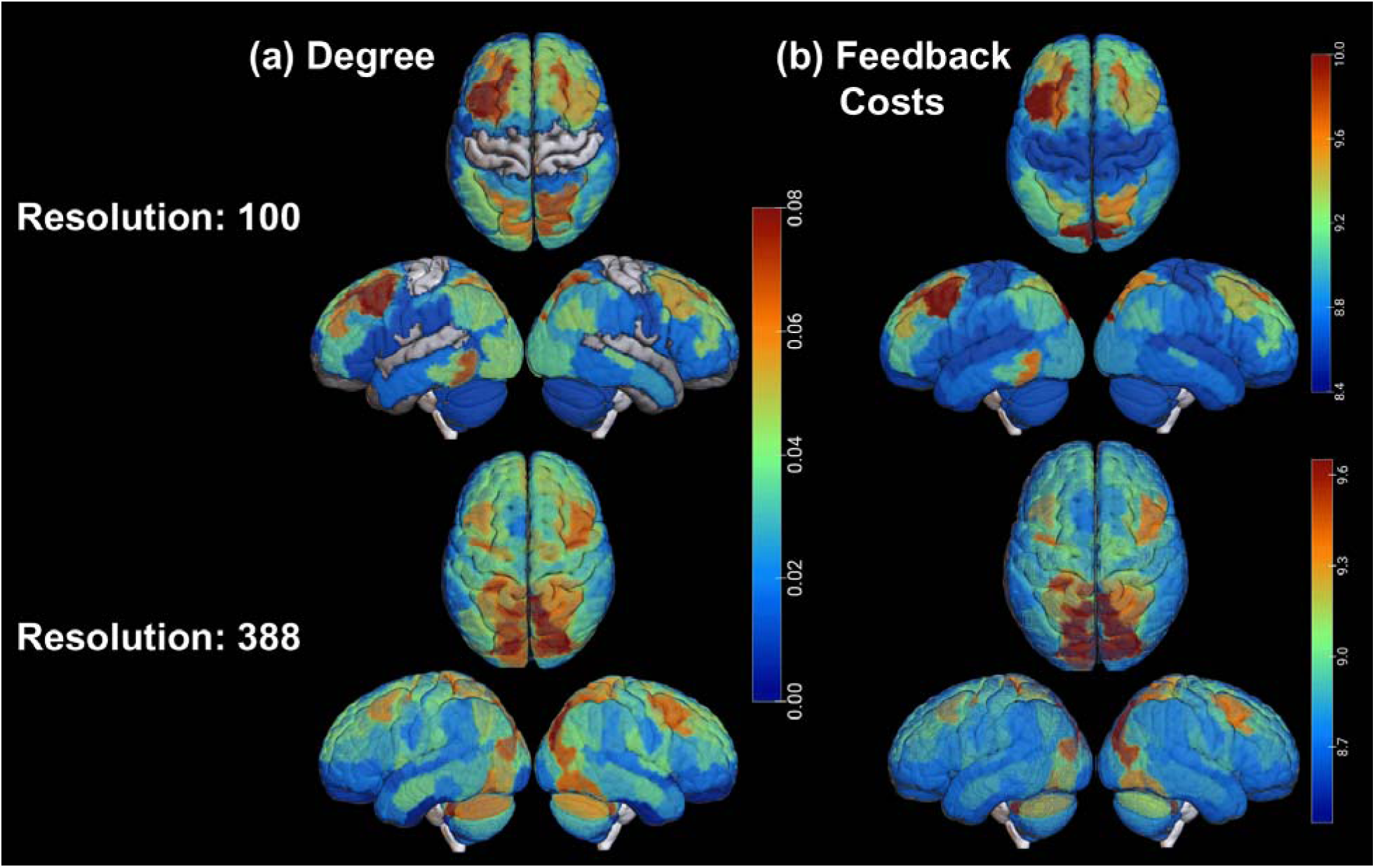
Spatial distributions of feedback costs (right panels) and regional connectedness (left panels) estimated at two spatial resolutions (100 and 388 regions) at baseline. In left panels, color corresponds to median (across the sample) node degree, normalized to the maximum possible degree (99 and 387, respectively). In right panels it corresponds to feedback costs at which regions lose their control action over the network.

### 3.2. Associations between feedback control costs and topological characteristics

The distributions of regional connectedness, frequency of intrinsic activation (dynamic region engagement), net information flow and feedback costs at which regions lost their control action, at baseline (top) and two-year follow-up (bottom), are shown in Figure 4. At both baseline and follow-up, similarity between the spatial distributions of degree and high control costs was high (median (IQR) BU: 0.91 (0.08) - 0.95 (0.05)), but moderate for dynamic region engagement (intrinsic regional activation) and control costs (median (IQR) BU: 0.59 (0.24) - 0.66 (0.16)). Ordinary linear regression models investigated associations between regions with top 25% and 10% critical feedback costs, respectively, and their topological properties.

**Figure 4:**
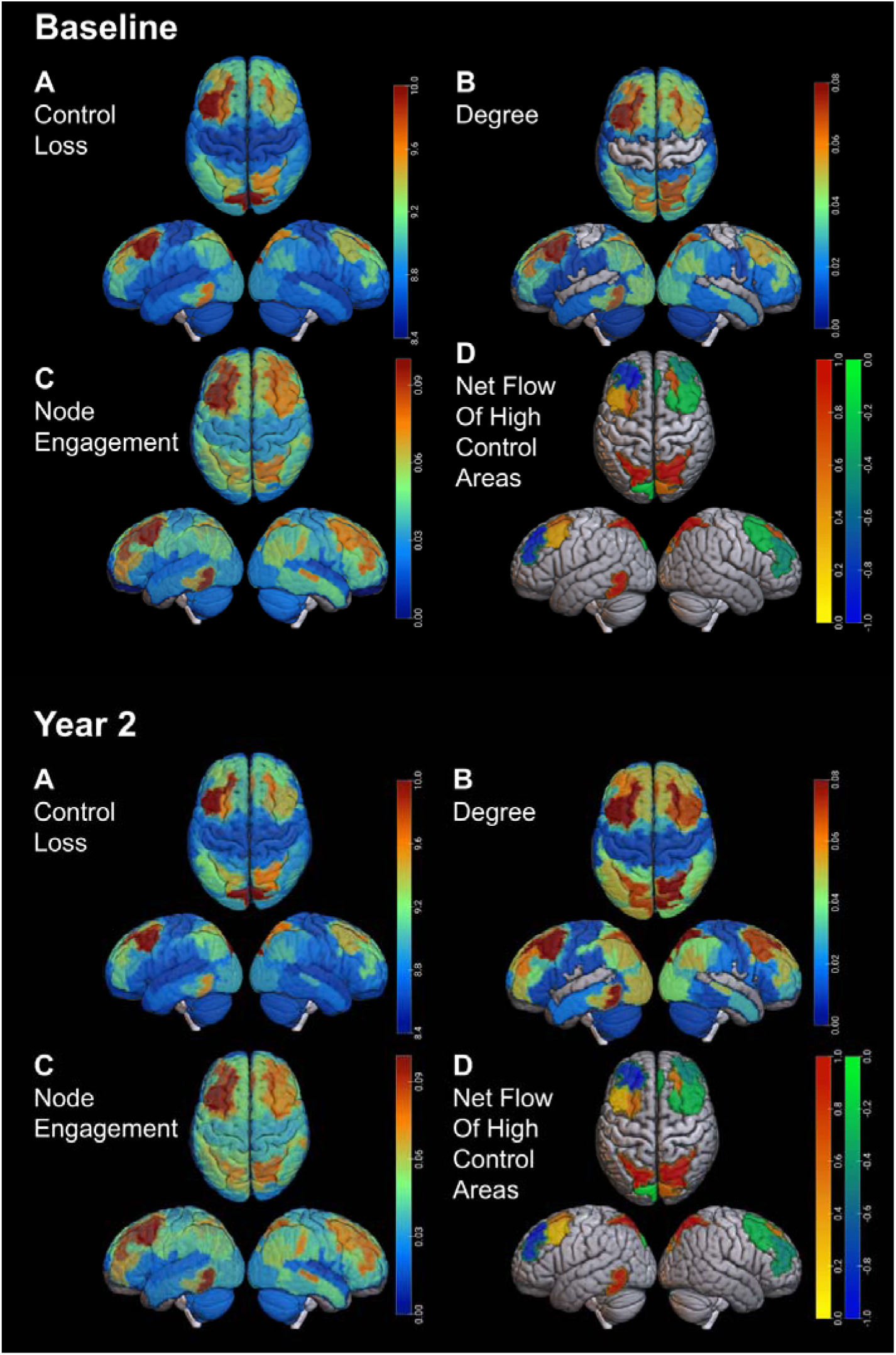
Distribution of control costs and topological properties estimated from the baseline (top panels) and two-year follow-up (bottom panels) data. (A): feedback costs associated with a region losing their control action over the network; (B): local connectedness (degree); (C): Frequency of node engagement; (D): net information flow, (positive net flow indicates that the region receives more information than it outputs).

#### A. Top 25% of costs

Regions consistently associated with the top 25% costs included posterior visual areas, bilateral prefrontal areas overlapping with frontoparietal control, salience and DM networks, bilateral parietal lobules overlapping with dorsal attention, control and default-mode networks, a left ventral posterior temporal area of the frontoparietal control network, and the right precuneus and retrosplenial cortex. Across assessments, degree was positively correlated with costs in all these regions, except for one area of the PFC (part of the DMN) at follow-up (baseline: p < 0.01, standardized regression coefficients (β) = 0.16 - 0.81, 95% confidence interval (CI) = [0.11, 0.85]); follow up: p < 0.01, β = 0.29 - 0.82, 95% CI = [0.23, 0.85]). Dynamic region engagement was positively correlated with feedback costs associated in most of these regions, both at baseline and follow up (baseline: p < 0.04, β = 0.06 to 0.24, 95% CI = [0.01, 0.29]; follow up: p < 0.01, β = 0.09 - 0.24, 95% CI = [0.04, 0.30]).

Across assessments, net information flow was negatively correlated with critical costs in multiple regions (i.e., regions associated with high feedback costs receive and output similar amounts of information and thus net flow is lower), including bilateral visual areas, prefrontal cortical areas overlapping with salience, control and default-mode networks, left inferior temporal gyrus, right precuneus and right retrosplenial cortex (p < 0.03, β = -0.33 to -0.06, 95% CI = [-0.38, -0.01]). At both assessments, net flow was positively associated with high feedback costs in the left ventral posterior temporal area (p < 0.01, β = 0.29, 95% CI = [0.23, 0.34] at baseline, and p < 0.01, β = 0.11 to 0.26, 95% CI = [0.05, 0.31] at follow up). Detailed associations and related model statistics are provided in Table 2.

**Table 1:**
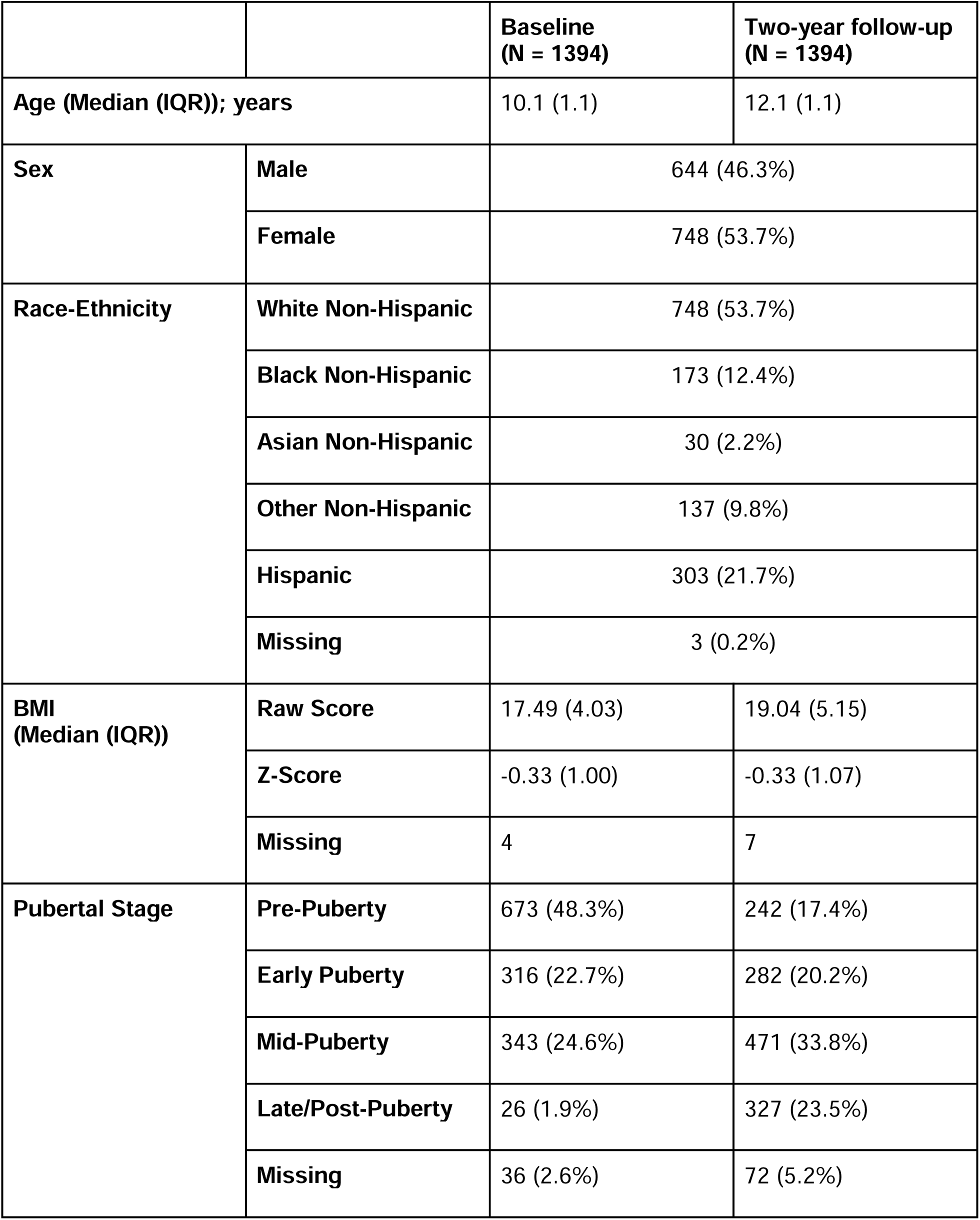
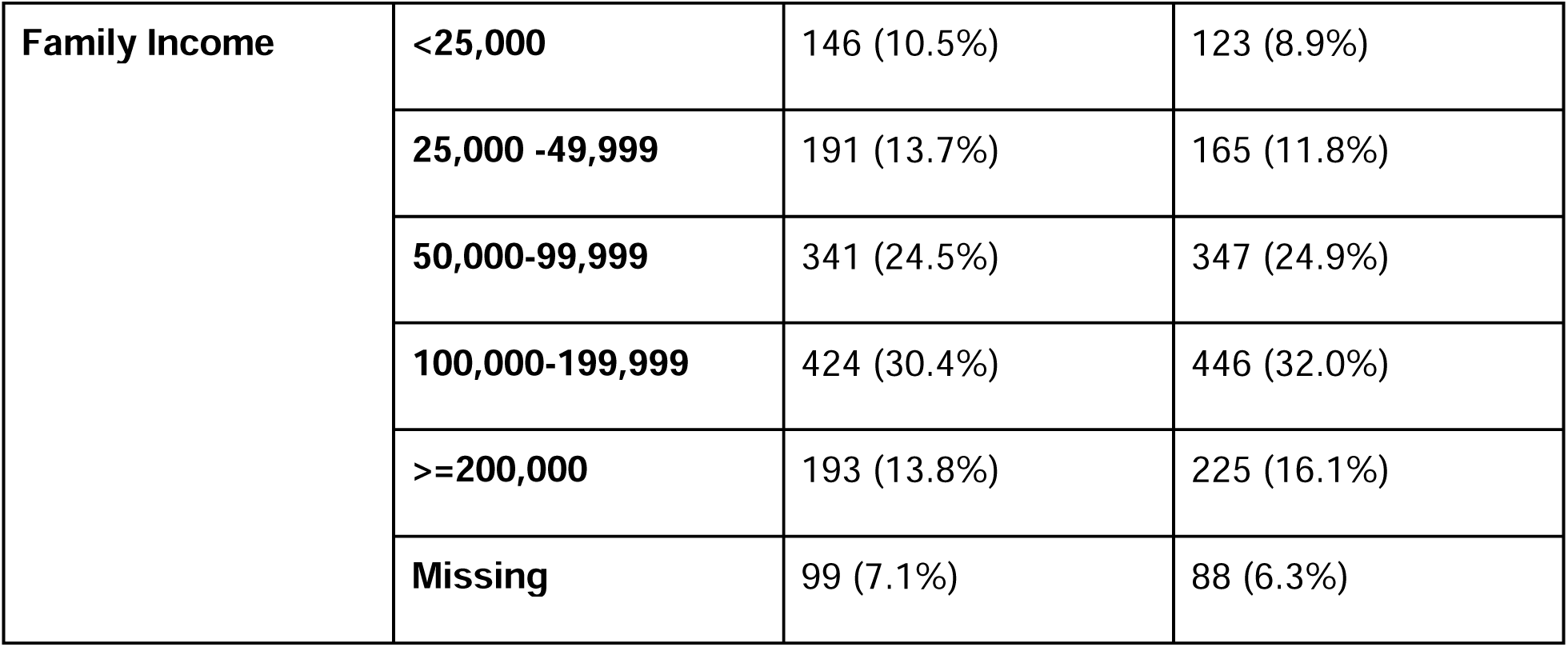
Participant characteristics at baseline and follow-up.

**Table 2:**
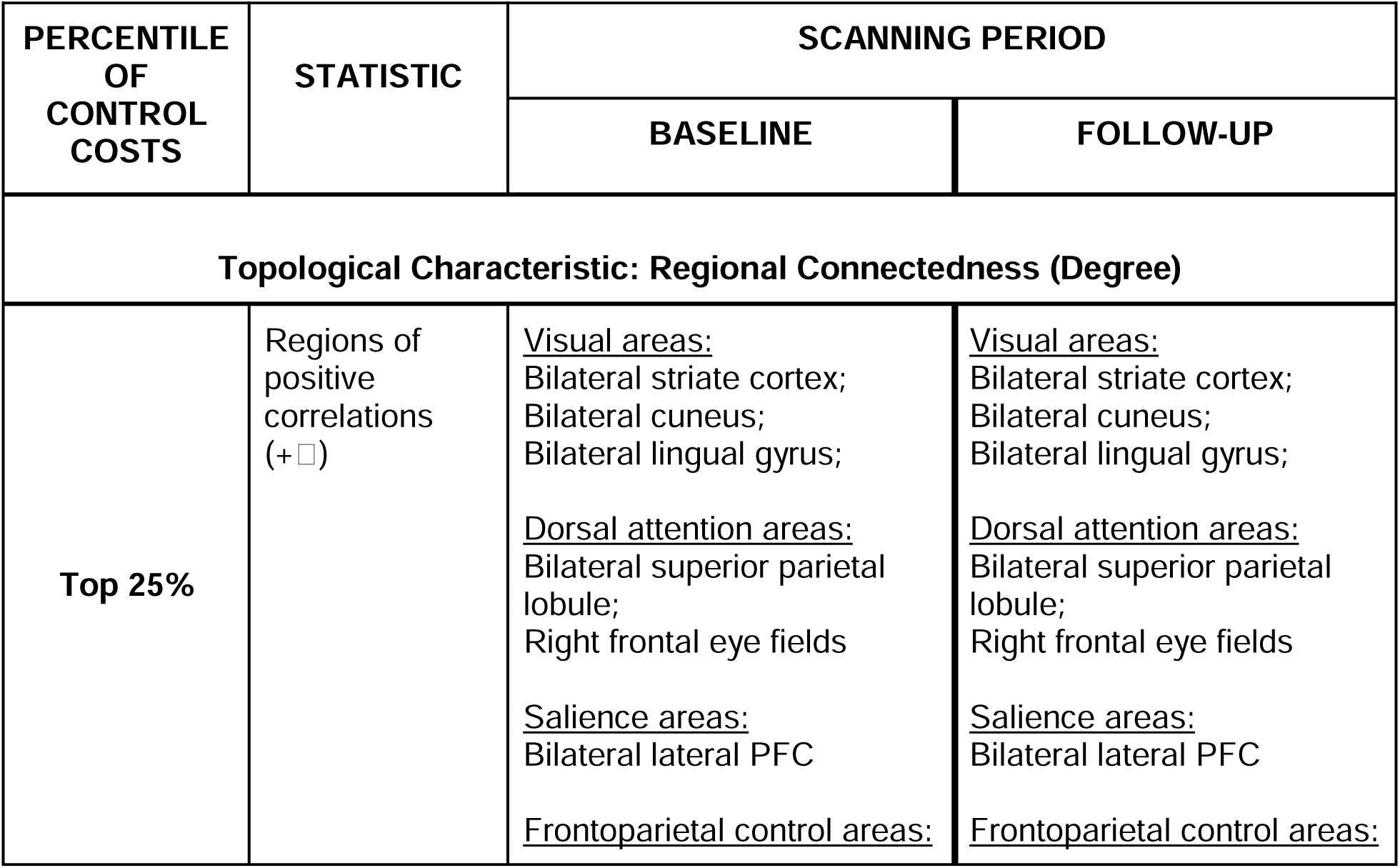
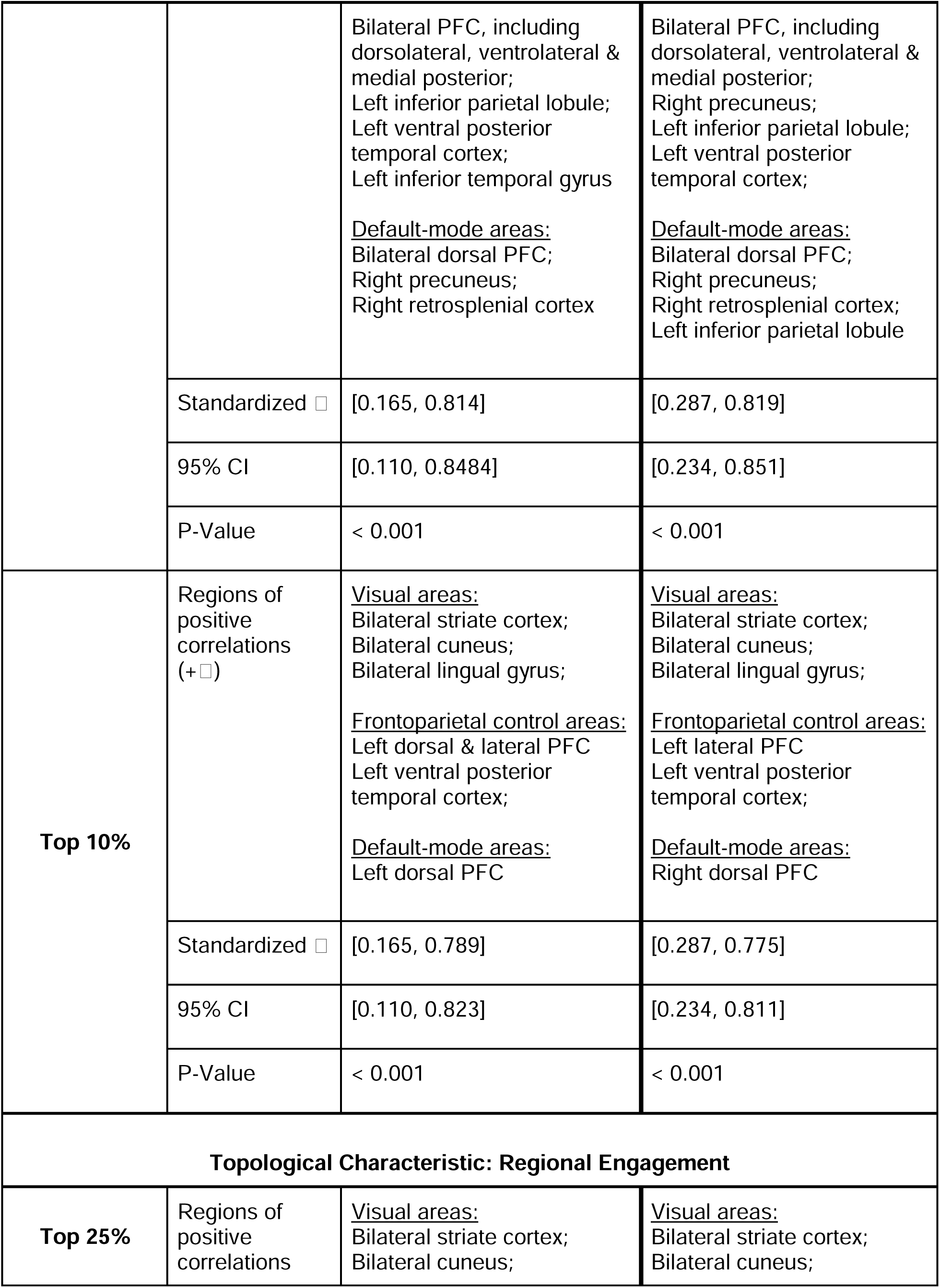
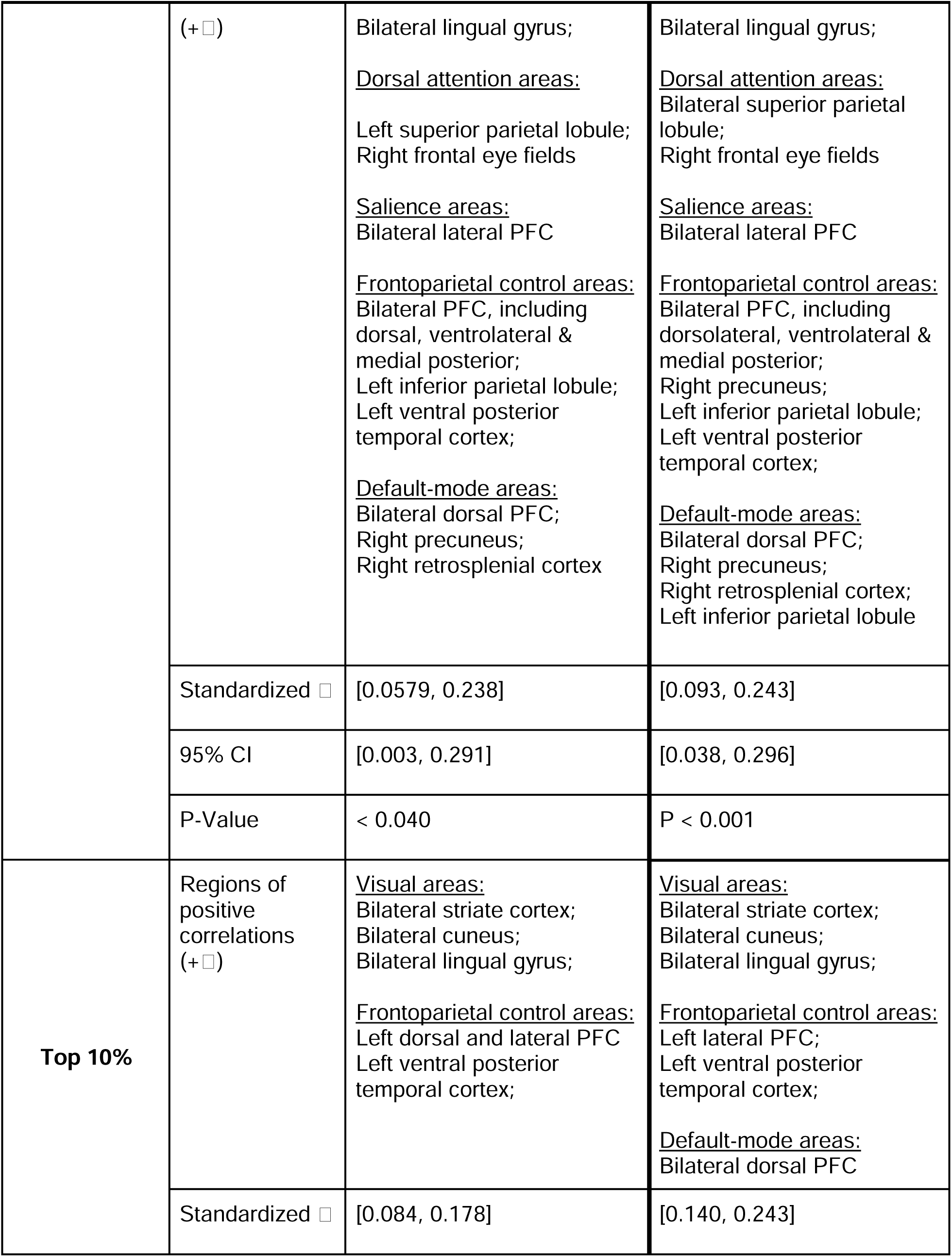
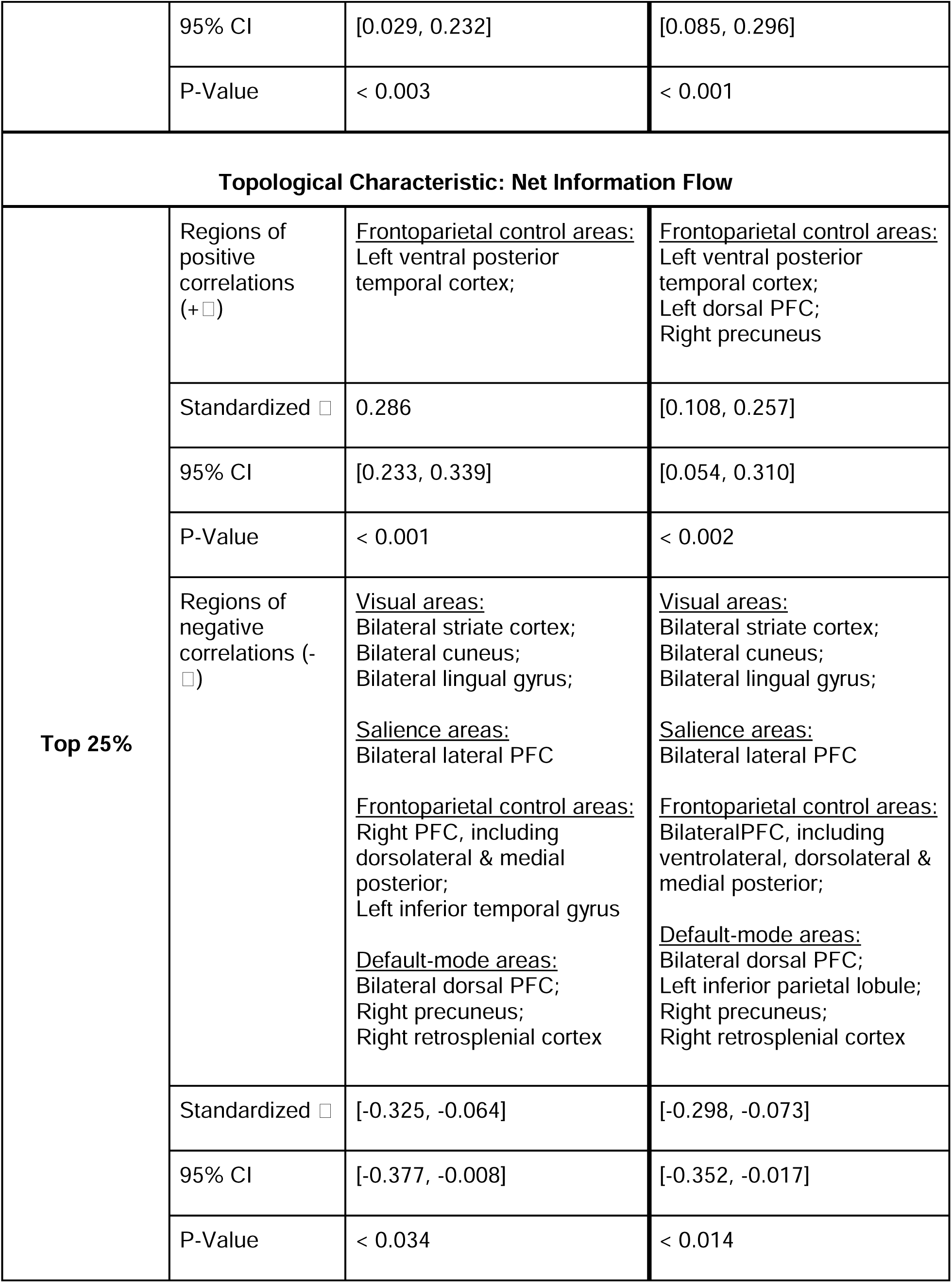
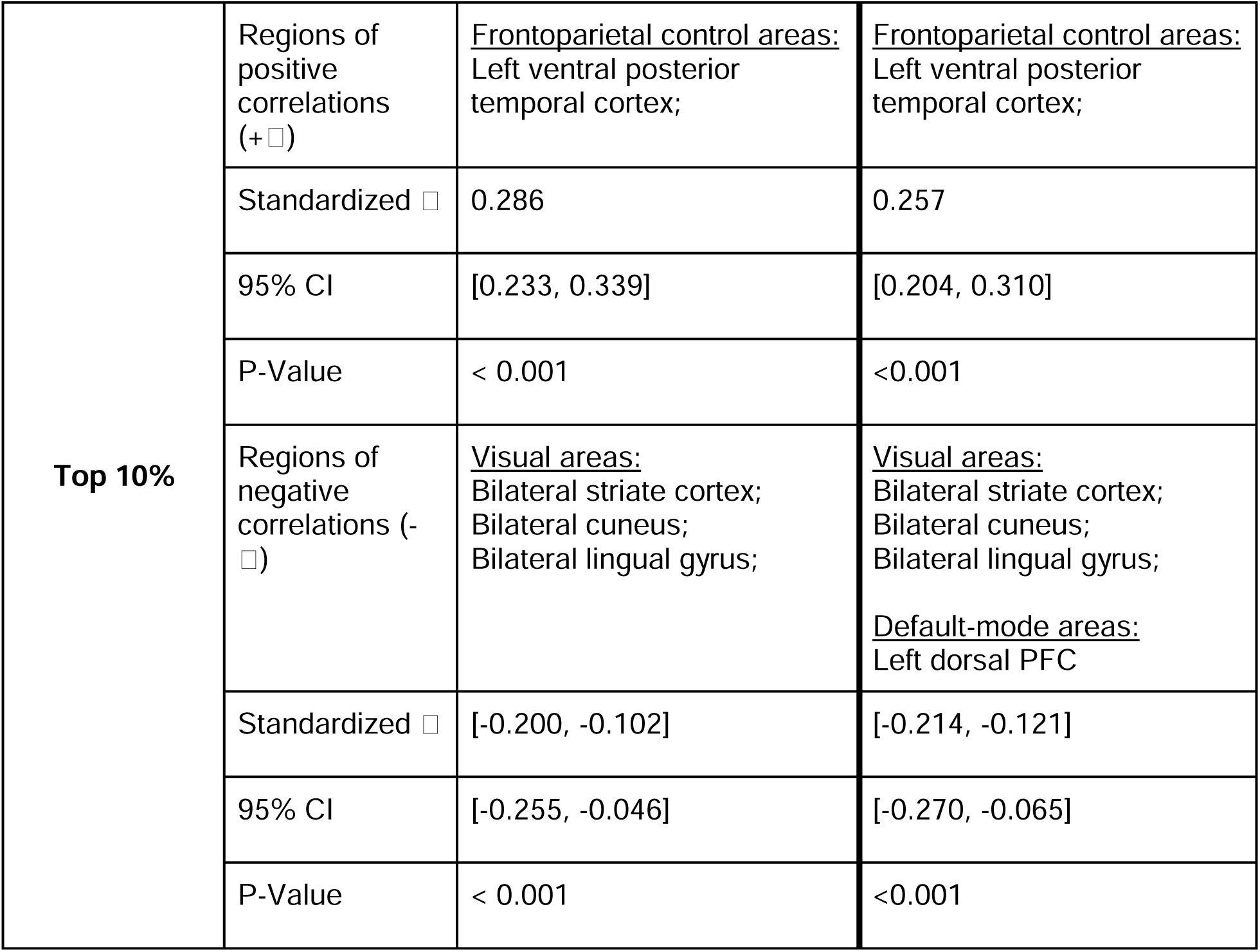
Statistics of ordinary linear regression models testing associations between control costs and topological characteristics. Reported p-values have been corrected for the False Discovery Rate

#### B. Top 10% of costs

Regions consistently associated with the top 10% highest feedback costs included bilateral posterior visual areas, left prefrontal and temporal areas that are part of the frontoparietal control network, and a left dorsal prefrontal region that is part of the DMN (the right dorsal PFC was also associated with high control costs at follow up). At both baseline and follow up, degree was positively correlated with feedback costs in all of these regions, with the exception of the dorsal prefrontal area at follow-up (baseline: p < 0.01, β = 0.16 to 0.79, 95% CI = [0.11, 0.82]; follow-up: p < 0.01, β = 0.29 to 0.78, 95% CI = [0.23, 0.81]). Dynamic engagement was positively correlated with feedback cost in most regions, with the exception of the dorsal prefrontal area at baseline (baseline: p < 0.01, β = 0.08 to 0.18, 95% CI = [0.03, 0.22]; follow-up: p < 0.01, β = 0.14 to 0.24, 95% CI = [0.09 to 0.30]).

At baseline, net information flow was negatively correlated with feedback costs in the bilateral posterior visual areas (p < 0.01, β = -0.20 to -0.10, 95% CI = [-0.25, -0.05]), and positively correlated in left ventral posterior temporal areas (p < 0.01, β = 0.29, 95% CI = [0.23, 0.34]). At follow-up, net flow was negatively correlated with feedback costs in the same regions, including left dorsal PFC (p < 0.01, β = -0.21 to -0.12, 95% CI = [- 0.27, -0.07]), but positively correlated in left ventral posterior temporal areas (p < 0.01, β = 0.26, 95% CI = [0.20, 0.31]). Detailed model statistics are provided in Table 2.

### 3.3 Pubertal changes in the distribution of feedback control costs

From baseline to follow-up, age-related differences in critical feedback costs were ≤1%.

Generalized linear mixed effects models that combined both baseline and follow-up samples, so that together they span the entire range of pubertal stages, were developed to examine changes in critical costs as a function of pubertal stage. Model statistics are summarized in Table 3. Consistently across runs, in most brain regions, puberty-related differences in critical feedback costs were nonsignificant. There were a few exceptions that were not consistent across runs. Based on data from the best (but not second) run, youth in later pubertal stages had higher feedback costs in the right cuneus (p = 0.01, β = 0.07, 95% CI = [0.02, 0.13]).

**Table 3:** Statistics of mixed effects linear regression models testing associations between feedback costs at which regions lose their control action over the network and pubertal stage in individual regions, separately for each analyzed runs. Reported p-values have been corrected for the False Discovery Rate (FDR).

| Associations Between Control Costs and Pubertal Stage |  |  |
| --- | --- | --- |
| Control Type | Statistic | Value |
| **BEST-QUALITY RUN** |  |  |
| **Cost at which regions lose control action over the network** | Regions of positive correlations (+$\beta$) | Right cuneus |
| | Standardized $\beta^*$ | 0.074 |
|  | 95% CI | [0.015, 0.133] |
|  | P-Value | 0.041 |
| **SECOND-BEST-QUALITY RUN** |  |  |
| None |  |  |

### 3.4 Associations between feedback costs and participant characteristics

Consistently across fMRI runs, girls had higher costs associated with self-control in the right ventral temporal area (p < 0.01, β = 0.01, 95% CI = [0.004, 0.016]) and lower costs in the left superior parietal lobule (p < 0.04, β = -0.011, 95% CI = [-0.019, -0.003]). They also had higher costs associated with loss of control action in the bilateral inferior temporal gyrus and the hippocampus (p < 0.05, β = 0.02 to 0.63, 95% CI = [0.001, 0.92]). They also had lower costs in the left superior parietal lobule and left postcentral gyrus (p = 0.02, β = -0.53 to -0.30, 95% CI = [-0.89, -0.08]). BMI was associated with higher self-control costs in the right visual lingual gyrus and cuneus (p < 0.03, β = [0.05, 0.06], 95% CI = [0.01, 0.10]), and higher costs associated with loss of control action in the left superior temporal gyrus (p < 0.02, β = 0.06, 95% CI = [0.02, 0.10]). There were no (consistent across runs) significant associations between critical control costs and race/ethnicity or family income. Model statistics are provided in Table 4. However, racial/ethnic minorities had lower critical control costs estimated from the best run in left prefrontal areas and the left inferior parietal cortex compared to the rest of the cohort (which is predominantly white and non-Hispanic; p < 0.04, β = -0.02 to -0.17, 95% CI = [-0.30, -0.04]). Model statistics are provided in Table S1.

**Table 4:**
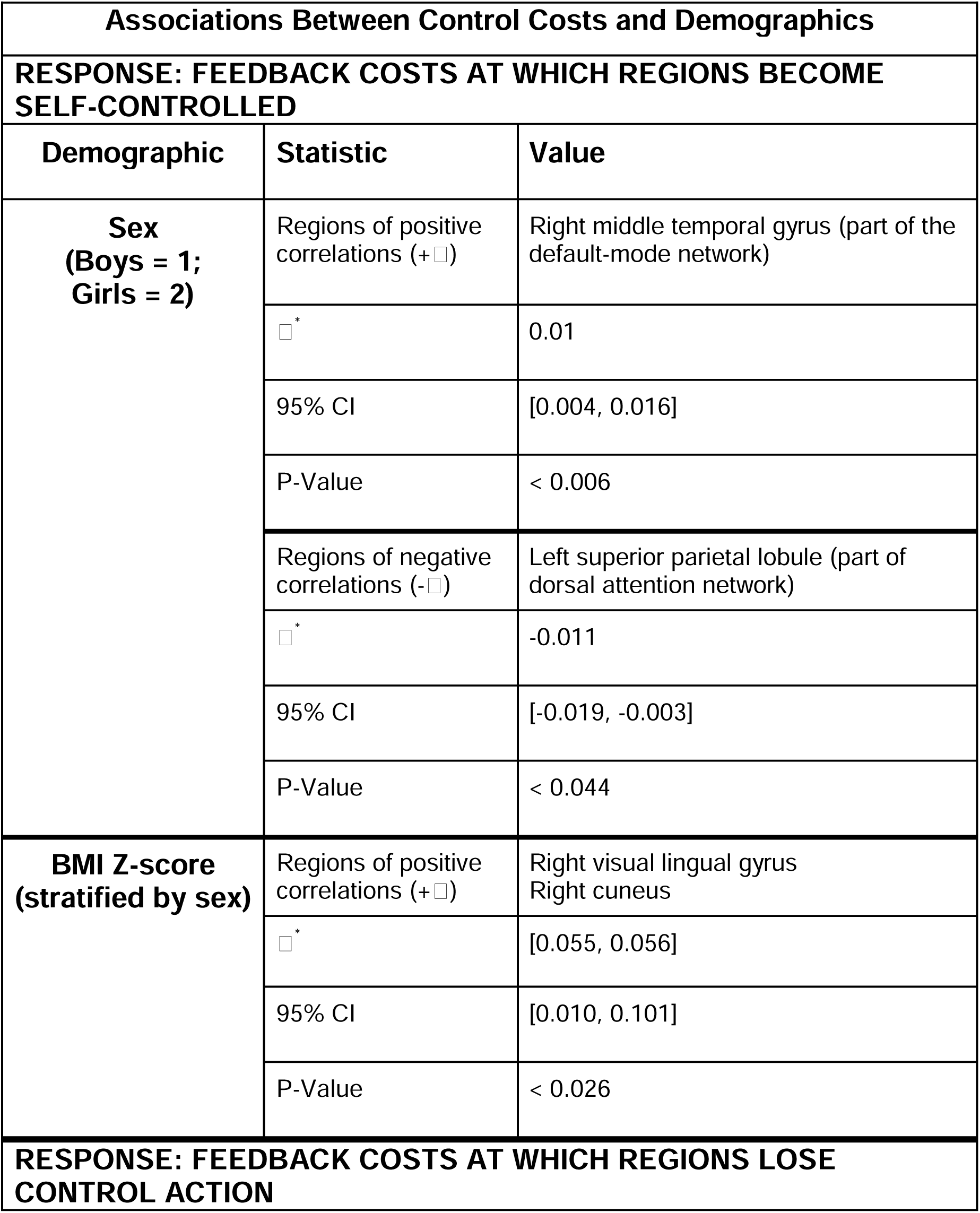
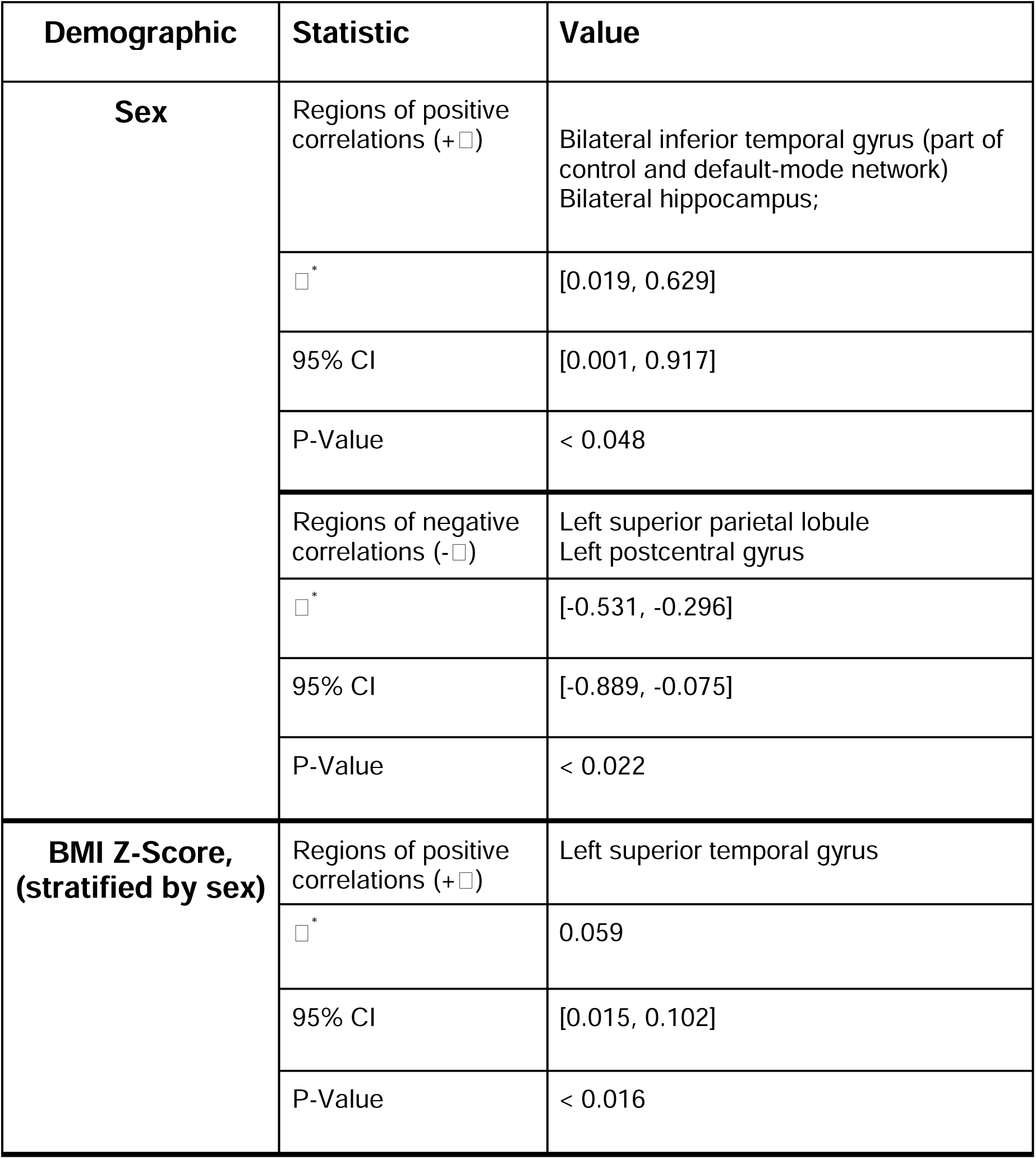
Statistics of mixed effects models testing associations between control costs and common demographics. Reported p-values have been corrected for the False Discovery Rate (FDR). Only statistics for regions that are repeated in both best and 2nd best run are reported.

## 3. DISCUSSION

In a large sample of youth with early longitudinal data from the ABCD study, we have used a novel to Neuroscience, sparsity-promoting feedback control framework to investigate the controllability of developing brain circuits, assuming that their dynamics and topology depend on internal (latent) mechanisms. These mechanisms may represent a combination of biological/biochemical processes that are not directly measurable, but their action on the organization of the developing circuitry can be estimated. We selected this framework, which also assumes that the controller’s action is sparse (i.e., a small number of regions with specific characteristics control the organization of the entire connectome), because it allowed us to incorporate the concepts of feedback penalty and sparsity, both critical to optimal network dynamics, and thus intrinsic and task-related synchronization of brain regions (Lizier et al, 2023). The adolescent brain is not fully developed, and thus its circuitry is not yet topologically or dynamically optimal. Tracking the controllability of the adolescent connectome may provide important novel insights into its developmental optimization. Two critical ranges of feedback costs were analyzed: a) costs at which the control action of individual regions only depends on themselves, and (typically high) costs at which regions lose their control action over the network. These costs were then examined as a function of regional topological properties, pubertal stage, and demographic characteristics.

In both regimes, high feedback costs were estimated in a small set of developed (primary visual) and developing (including prefrontal) regions. Higher costs were associated with higher regional connectedness and higher dynamic consistency, i.e., the region was consistently activated during spontaneous brain coordination at rest. Posterior areas associated with high feedback costs included regions that are part of the peripheral visual network and are well developed in adolescence, but also retrosplenial cortex, the cuneus and the superior parietal lobule (SPL), all of which continue to develop during adolescence as visual processing is refined, but also support other cognitive processes that evolve during this period (Vann et al, 2009; Miller et al, 2014; Wang et al, 2015; Mitchell et al, 2018; Palejwala et al, 2021). Other areas associated with high feedback costs were parts of the ventral temporal cortex (involved in visual recognition and categorization), which also continues to maturate in adolescence (Conway, 2018; Nordt et al, 2023). Together with the cuneus, SPL, and retrosplenial cortex, these areas are involved in information integration and support multiple cognitive processes, including somatomotor processing, spatial navigation and shifting, memory, attentional control, but also posture and body part localization (Caminiti et al, 1996; Wopert et al, 1998; Wolbers et al, 2003; Makino et al, 2004; Felician et al, 2004; Molenberghs et al, 2007; Koenigs et al, 2009; Vandenberghe et al, 2011; Schmostein, 2012; Lester et al, 2014; Passarelli et al, 2021; Balcerek et al, 2021) and are, therefore, cognitive hubs.

A number of regions associated with high feedback costs were in the prefrontal cortex, which undergoes accelerated maturation during adolescence, as high-level cognitive processes continue to develop (Nelson and Guyer, 2011; Somerville et al, 2013; Dumontheil, 2014; Caballero et al, 2016; Kolk and Rakic, 2022). These regions included dorsal and lateral prefrontal cortices, which support executive function and decision-making, memory and learning, and cognitive control (Wagner et al, 2001; Petrides, 2005; Fuster, 2015; Nee and D’Esposito, 2016), and are cognitive hubs as well (Gray et al, 2002; Milan et al, 2016; Menon and D’Esposito, 2022; Friedman and Robbins, 2022). Furthermore, together with the posterior areas associated with high feedback costs, they are part of a core set of brain regions that play a ubiquitous role in cognitive function (Dosenbach et al, 2006; Assem et al, 2020).

Most of the regions associated with high feedback costs have also been identified as topological hubs (i.e., highly connected and central to the organization of the connectome) that continue to evolve in adolescence (Tomasi and Volkow, 2011; Park and Friston, 2013; van der Heuvel and Sporns, 2013; Power et al, 2013; Hwang et al, 2013; Bertolero et al, 2015; Baker et al, 2015; Gordon et al, 2018; Oldham and Fornito, 2019). An independent investigation of these regions’ topological characteristics in a sample from the ABCD study that overlapped with this study’s sample has classified several of them as hubs that are stable during puberty (Gibble et al, 2024 (unpublished manuscript)). Higher feedback costs were associated with higher regional connectedness, suggesting that hubs maintain their control action over the connectome even at high costs. In addition, they were dynamically consistent, i.e., they were repeatedly highly connected during spontaneous synchronization of brain regions. Finally, in most of these regions (with the exception of ventral temporal cortex) higher feedback costs were associated with lower net information flow through them, i.e., with lower differences between information input and output from these regions. Brain hubs play a critical role in the communication between spatially distributed brain regions, by integrating information from them (e.g., resulting from domain-specific computations), and distributing the output of this synthesis in response to cognitive demands (van den Heuvel et al, 2012). Thus, information influx and outflux may be approximately equal in these regions, and thus net flow is low. The correlation between high feedback costs and low net flow again indicates that the identified regions may indeed be brain hubs, and suggests a positive association between controllability and communication cost.

Critical control costs were highly reproducible across thresholds, fMRI runs (and thus snapshots of brain activity) and assessments, i.e., from ages 9 - 12 years. Furthermore, in most regions (with the exception of the right cuneus) these costs did not significantly change with pubertal stage, suggesting that their control action may be invariant to developmental changes, even if their local topology changes during this period. Indeed, many of these regions are elements of the DM and frontoparietal control networks, which are partly inter-connected (Dixon et al, 2018), and their topological organization and properties continue to develop significantly in adolescence (Fair et al, 2008; Sherman et al, 2014).

Despite its many strengths, including a novel to the field control framework, large early longitudinal sample that captured the heterogeneity of the developing adolescent brain, and cutting-edge analytic approaches for estimating topological/dynamic characteristics, the study had some limitations. Although the sample spanned the entire range of pubertal stages, it covered ages ∼9-12 years, and not the entire adolescent period. To date, there are no other equally large samples covering this period. However, as the ABCD study continues to follow brain development in this cohort, additional longitudinal data will become available. Thus, a future study could extend this investigation and track the brain’s controllability, and the identified regions associated with high feedback costs, at later ages. Furthermore, this study purposely focused only on the resting-state connectome, and estimated static and dynamic topological characteristics and information flow, directly from the fMRI time series. A few prior studies have examined the controllability of the structural connectome, but such an investigation was outside the scope of the present study. A future study could compare the controllability of structural and functional circuits in the same cohort, since the ABCD also collects diffusion MRI data. Finally, the control problem was solved assuming linear system dynamics, which may be a simplification of potentially much more complex brain dynamics. However, a recent study in adults has shown that at the macroscale, brain dynamics are well described by linear models (Nozari et al, 2024), although it is unclear whether this is also the case in the developing brain.

Despite a few limitations, this study makes a significant contribution to the field by addressing the fundamental question of how network dynamics are controlled in the developing brain, which is suboptimally organized and changes significantly, especially in adolescence. To answer this question, the study assumed a closed-loop control framework (which by design uses internal feedback as a control mechanism), and imposed a sparsity constraint, which is biologically meaningful given that brain network dynamics are likely controlled by a small set of regions with specific characteristics. To the best of our knowledge, this is the first large-scale application of this framework to developing brain networks. The study has identified a parsimonious set of highly reproducible (across snapshots of resting-state activity) and developmentally stable regions that exert their action on intrinsic brain network dynamics even at high feedback costs. Most of these regions are core cognitive hubs and highly connected functional hubs that are central to the organization of the connectome and cognitive function across domains. These results provide novel insights into an important yet relatively underexplored mechanism of brain network dynamics. They suggest that even the underdeveloped (and in some areas redundantly connected) adolescent brain, fundamental mechanisms of system controllability may be well developed, as reflected in the small set of distributed regions that exert their action on the connectome, to facilitate information processing and response to cognitive demands.

## Data and code availability statements

The data analyzed in this study are publicly available through the National Institute of Mental Health Data Archive (NDA): https://nda.nih.gov/.

The accession number (DOI) associated with study in NDA repository is: http://dx.doi.org/10.15154/1523041

## Codes used in the data analyses are available at

- https://github.com/cstamoulis1/Next-Generation-Neural-Data-Analysis-NGNDA/
- https://github.com/cstamoulis1/Closed-Loop-Control-of-Developing-Brain-Networks/

## Ethics statement

This study involved secondary analyses of publicly available, deidentified human data and was approved by the Institutional Review Board.

## SUPPLEMENTAL MATERIALS

**Figure S1:**
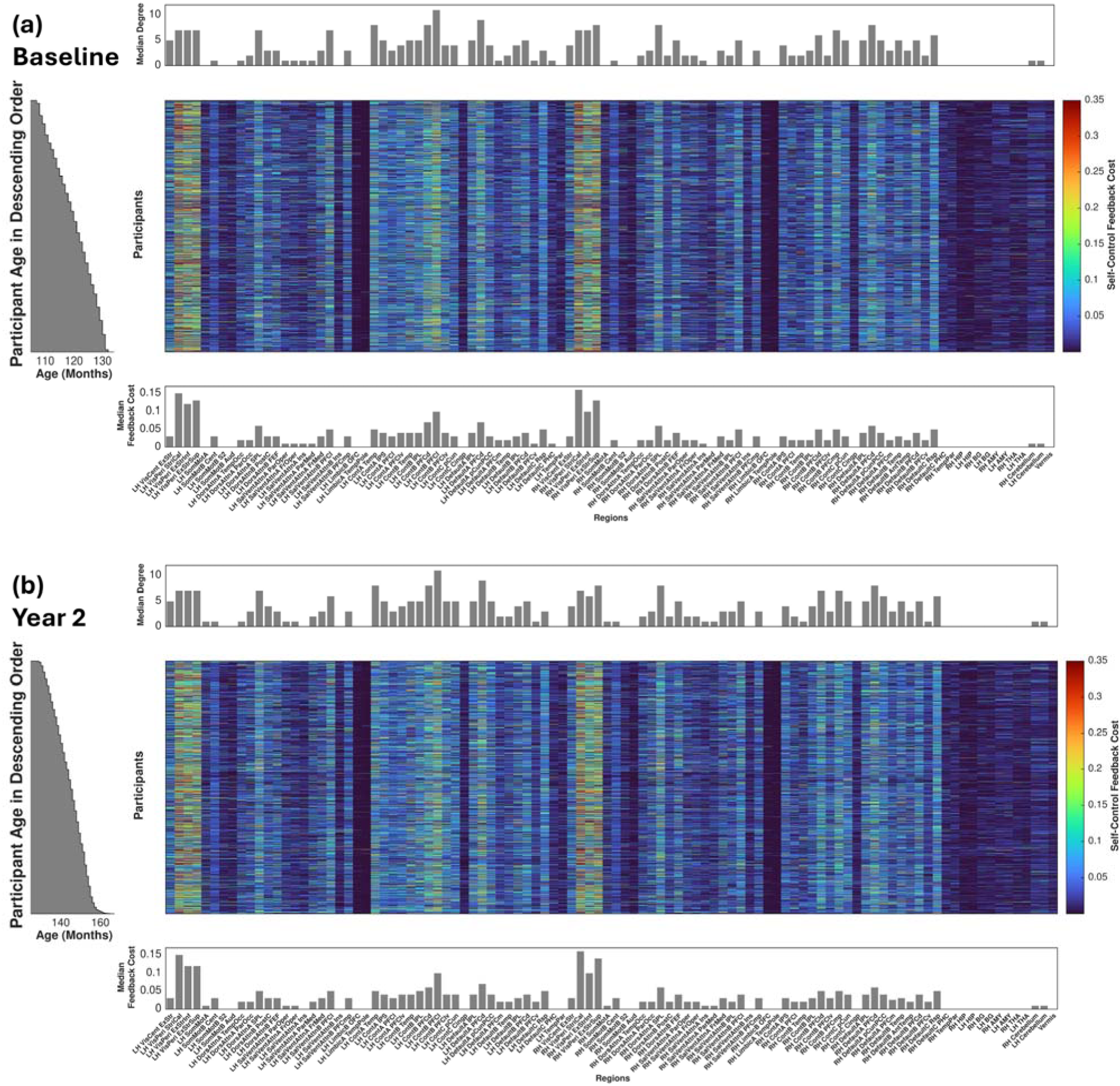
Feedback costs at which a region’s control action only depends on its own dynamic state, as a function of participants (y-axis) and analyzed regions (x-axis). The color range corresponds to cost values, estimated from the best-quality run at baseline (a) and two-year follow up (b). Participants are sorted by age. Median (over participants) regional connectedness (degree) for each analyzed region is shown in the bar plot above each heat map, and median feedback costs in the bar plot below each heat map. All parameters were estimated from adjacency matrices obtained by thresholding corresponding connectivity matrices based on the moderate outlying peak cross-correlation value.

**Figure S2:**
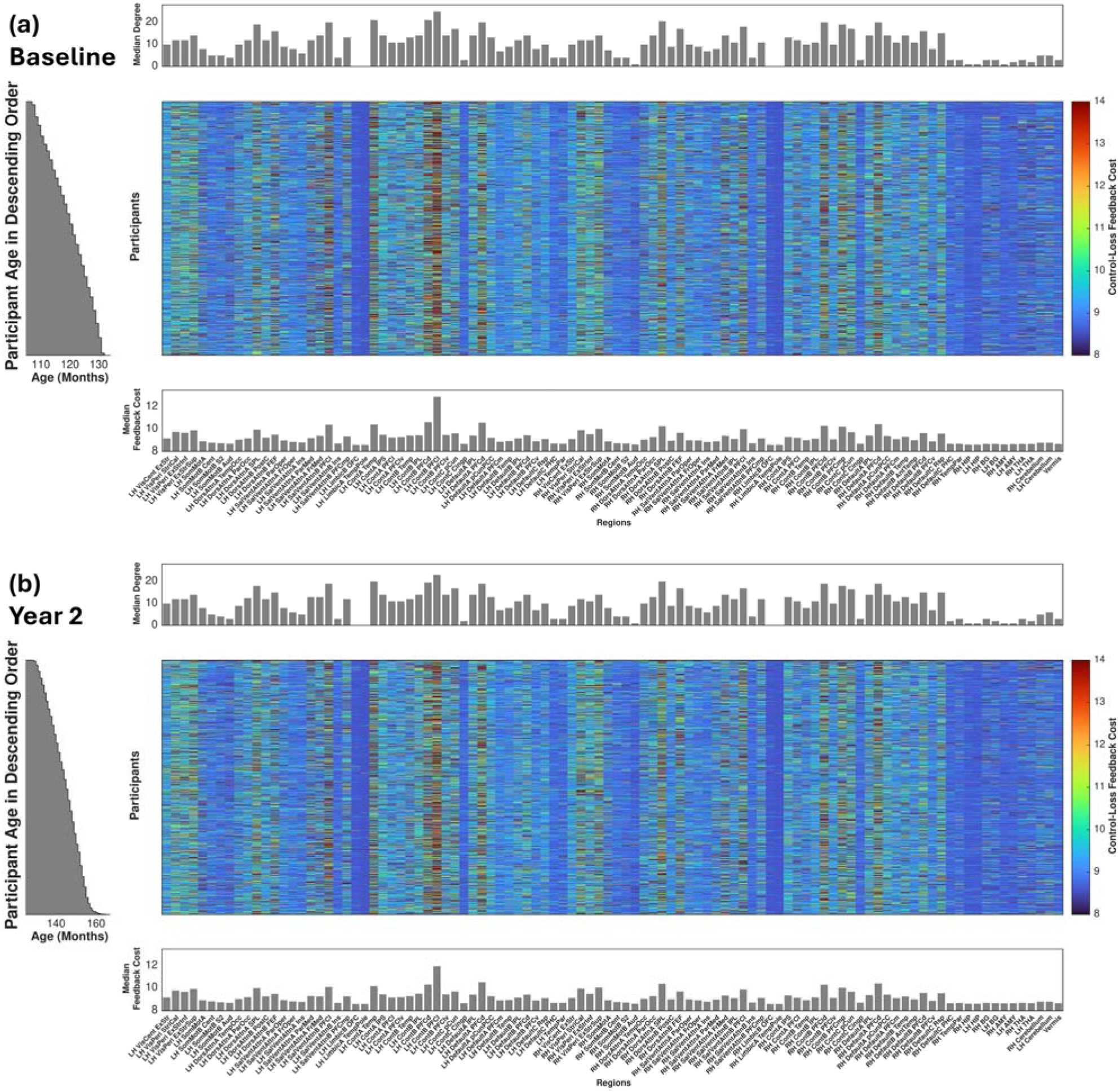
Feedback costs at which regions lose their control action on the network, as a function of participants (y-axis) and analyzed regions (x-axis). The color range corresponds to cost values, estimated from the best-quality run at baseline (a) and two-year follow up (b). Participants are sorted by age. Median (over participants) regional connectedness (degree) for each analyzed region is shown in the bar plot above each heat map, and median feedback costs in the bar plot below each heat map. All parameters were estimated from adjacency matrices obtained by thresholding corresponding connectivity matrices based on the 75^th^ percentile of peak cross-correlation value.

**Figure S3:**
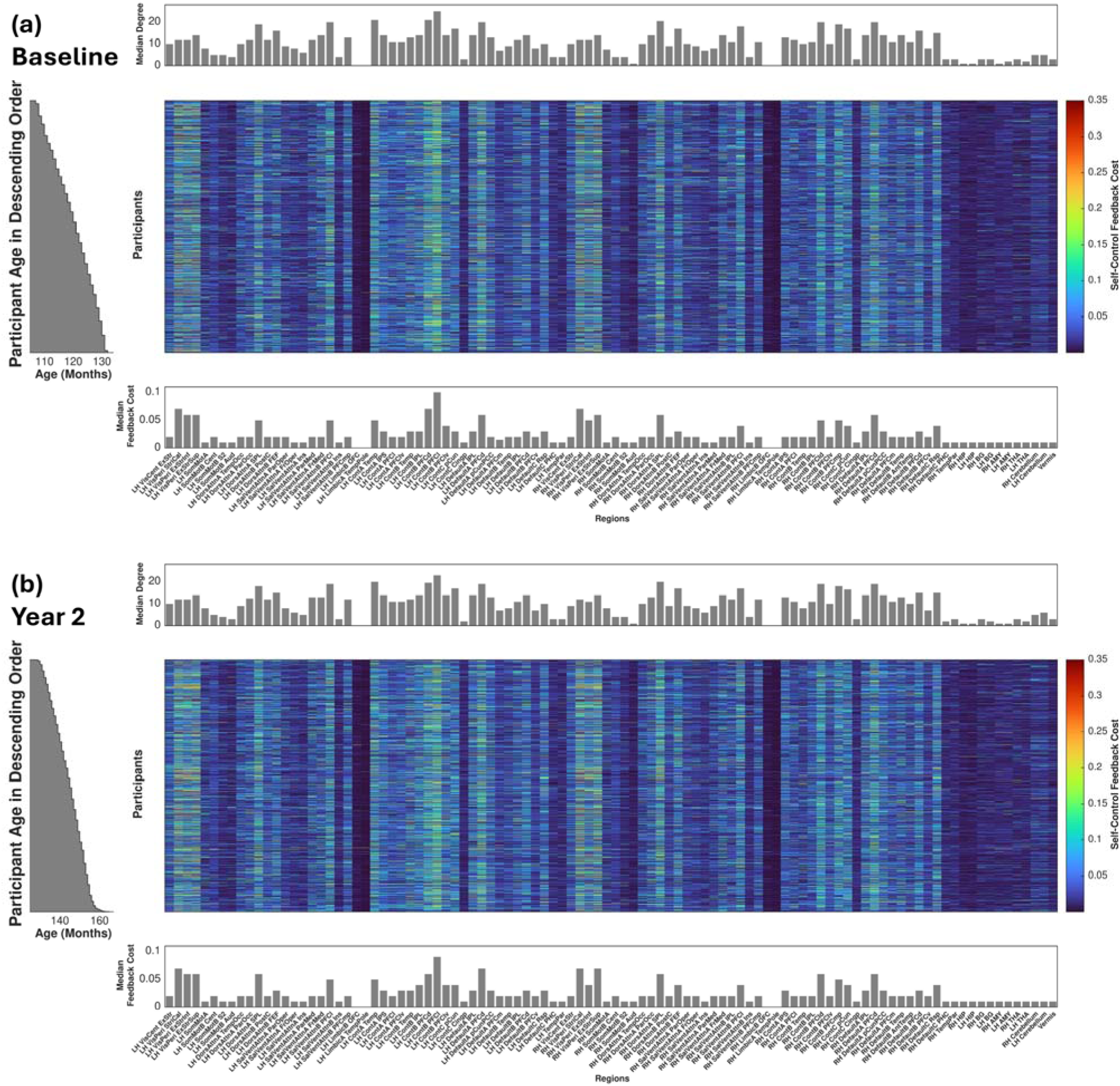
Feedback costs at which a region’s control action only depends on its own dynamic state, as a function of participants (y-axis) and analyzed regions (x-axis). The color range corresponds to cost values, estimated from the best-quality run at baseline (a) and two-year follow up (b). Participants are sorted by age. Median (over participants) regional connectedness (degree) for each analyzed region is shown in the bar plot above each heat map, and median feedback costs in the bar plot below each heat map. All parameters were estimated from adjacency matrices obtained by thresholding corresponding connectivity matrices based on the 75^th^ percentile of peak cross-correlation value.

**Figure S4:**
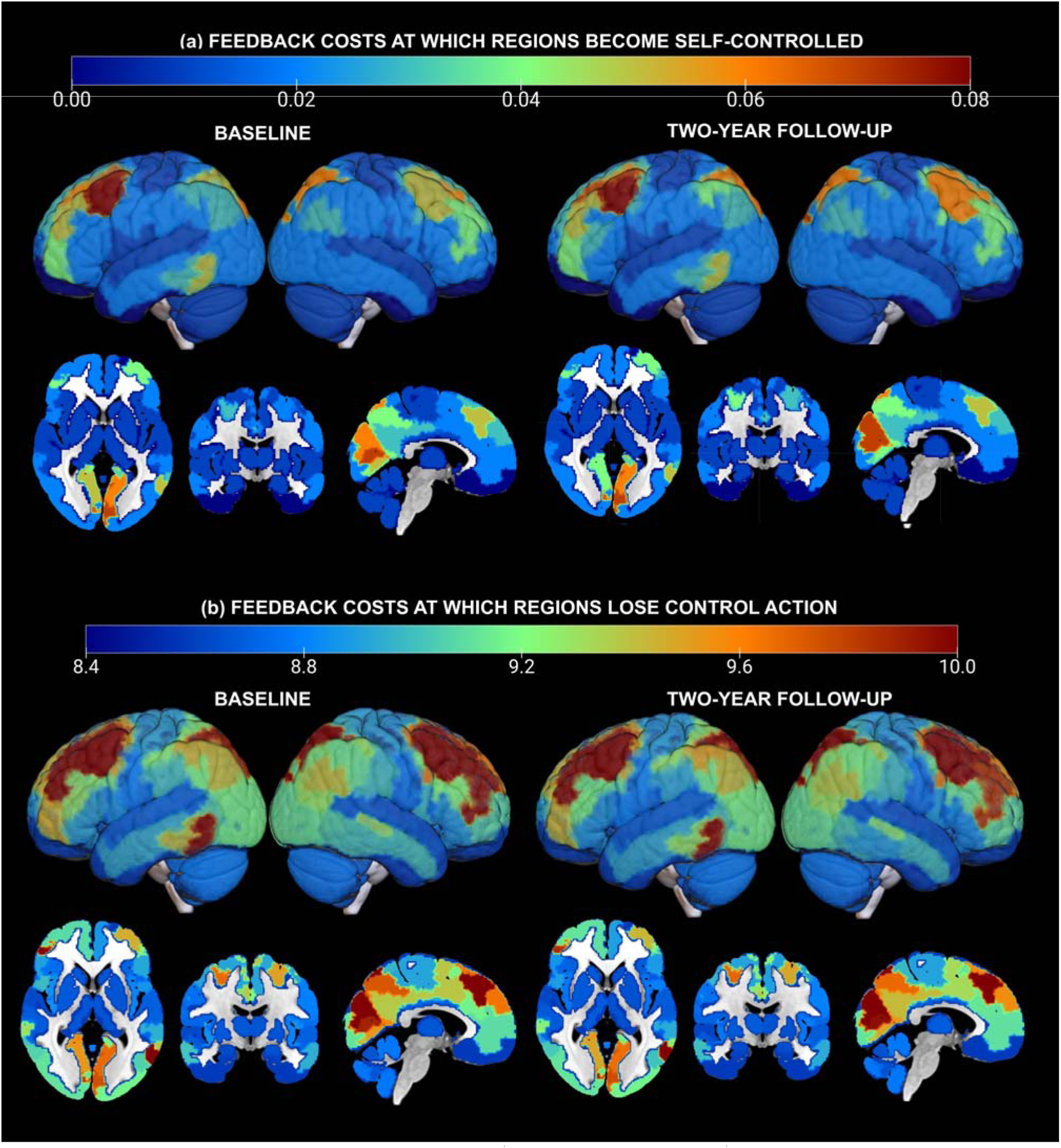
Spatial distribution of median (across participants) feedback costs at which (a) a region’s control action depends only on its own dynamic state, and (b) a region loses its control action over the network. Distributions are shown separately for baseline (left) and two-year follow up (right), and two-dimensional (horizontal, coronal, sagittal) slices and three-dimensional volumes are superimposed. Colors correspond to feedback cost values. All parameters were estimated from adjacency matrices obtained by thresholding the corresponding connectivity matrices based on the 75th peak cross-correlation value.

**Figure S5:**
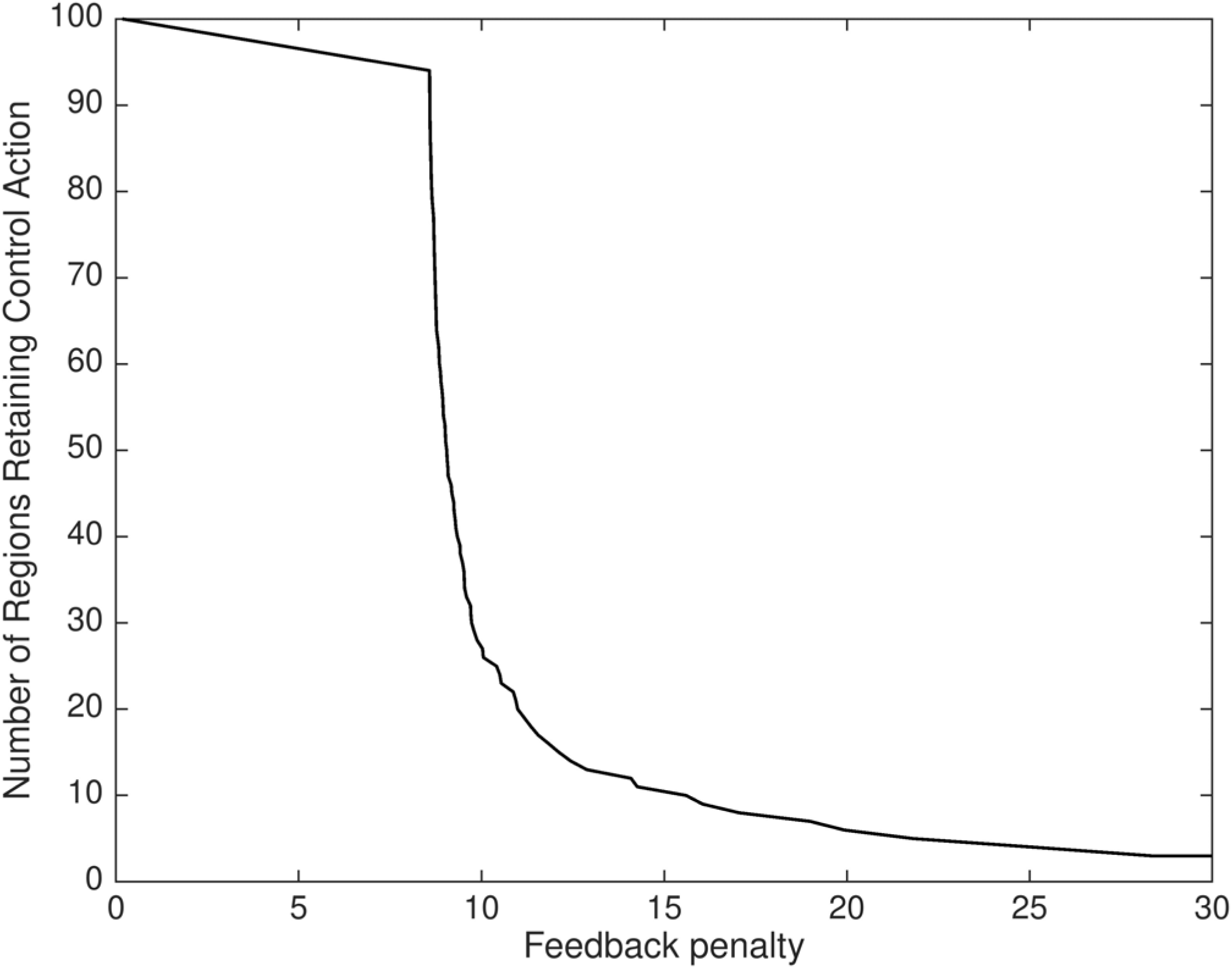
Example of number of regions that retain their control action as a function of increasing feedback cost (data are based on a representative brain from the analyzed sample).

**Table S1:**
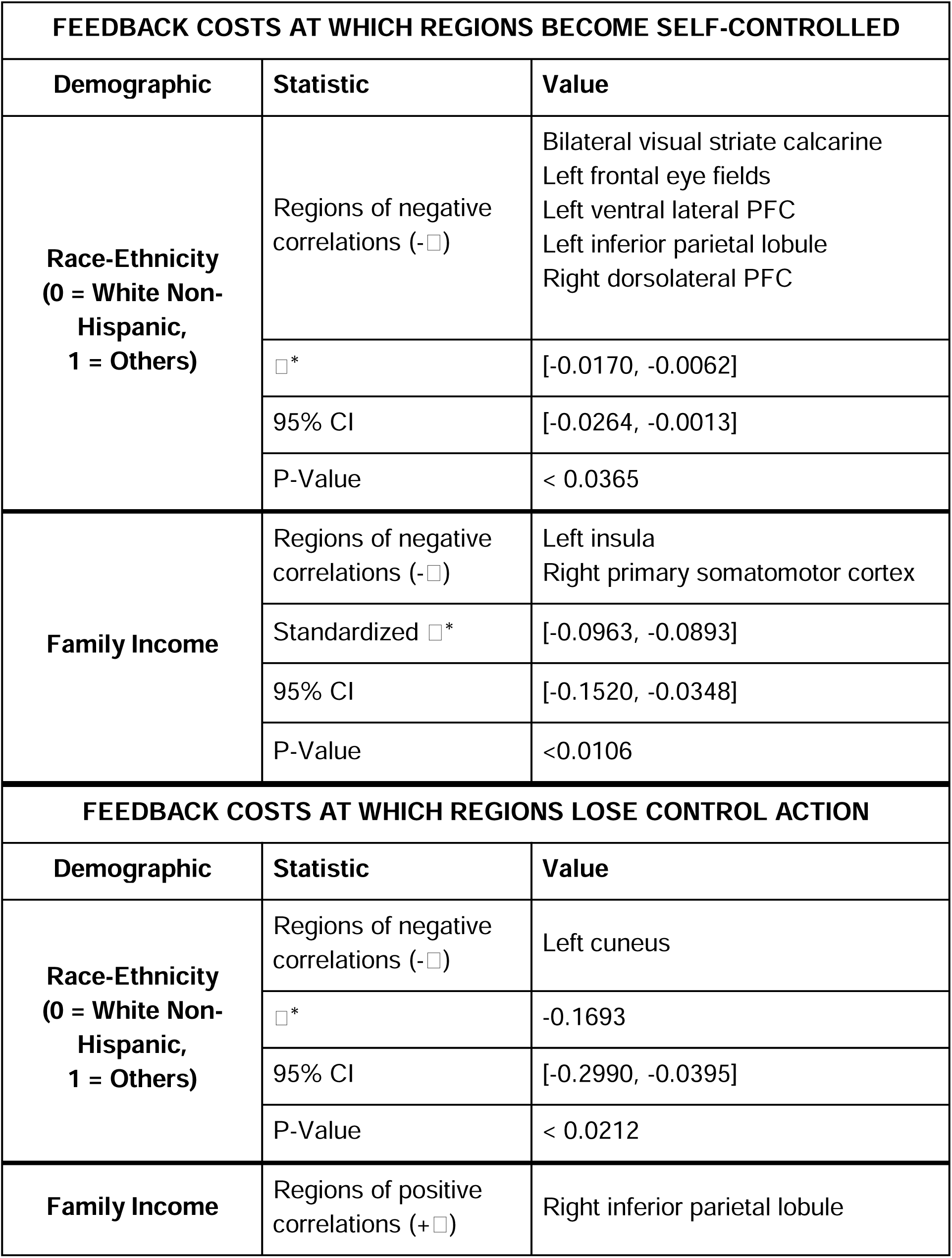
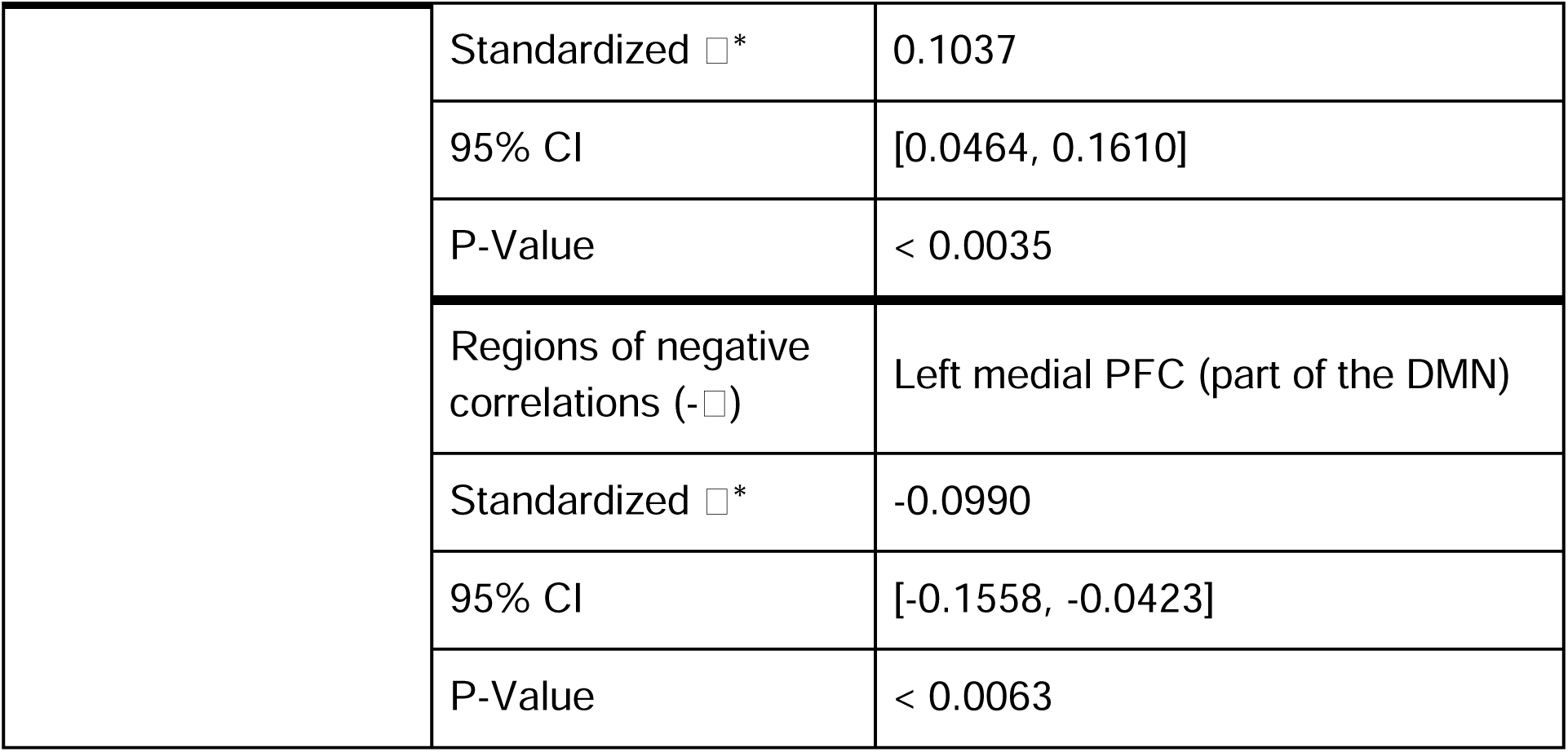
Statistics of mixed effects models testing associations between control costs (estimated from the best-quality fMRI run) and demographic characteristics. Reported p-values have been corrected for the False Discovery Rate (FDR).

| FEEDBACK COSTS AT WHICH REGIONS BECOME SELF-CONTROLLED |  |  |
| --- | --- | --- |
| Demographic | Statistic | Value |
| Race-Ethnicity<br>(0 = White Non-Hispanic,<br>1 = Others) | Regions of negative correlations ($-\beta$) | Bilateral visual striate calcarine<br>Left frontal eye fields<br>Left ventral lateral PFC<br>Left inferior parietal lobule<br>Right dorsolateral PFC |
| | $\beta^*$ | [-0.0170, -0.0062] |
|  | 95% CI | [-0.0264, -0.0013] |
|  | P-Value | < 0.0365 |
| Family Income | Regions of negative correlations ($-\beta$) | Left insula<br>Right primary somatomotor cortex |
| | Standardized $\beta^*$ | [-0.0963, -0.0893] |
|  | 95% CI | [-0.1520, -0.0348] |
|  | P-Value | <0.0106 |
| FEEDBACK COSTS AT WHICH REGIONS LOSE CONTROL ACTION |  |  |
| Demographic | Statistic | Value |
| Race-Ethnicity<br>(0 = White Non-Hispanic,<br>1 = Others) | Regions of negative correlations ($-\beta$) | Left cuneus |
| | $\beta^*$ | -0.1693 |
|  | 95% CI | [-0.2990, -0.0395] |
|  | P-Value | < 0.0212 |
| Family Income | Regions of positive correlations ($+\beta$) | Right inferior parietal lobule |
| | Standardized $\beta^*$ | 0.1037 |
|  | 95% CI | [0.0464, 0.1610] |
|  | P-Value | < 0.0035 |
| | Regions of negative correlations ($-\beta$) | Left medial PFC (part of the DMN) |
| | Standardized $\beta^*$ | -0.0990 |
|  | 95% CI | [-0.1558, -0.0423] |
|  | P-Value | < 0.0063 |

