## Supplemental materials for "The Adolescent Functional Connectome is Dynamically Controlled by a Sparse Core of Cognitive and Topological Hubs"

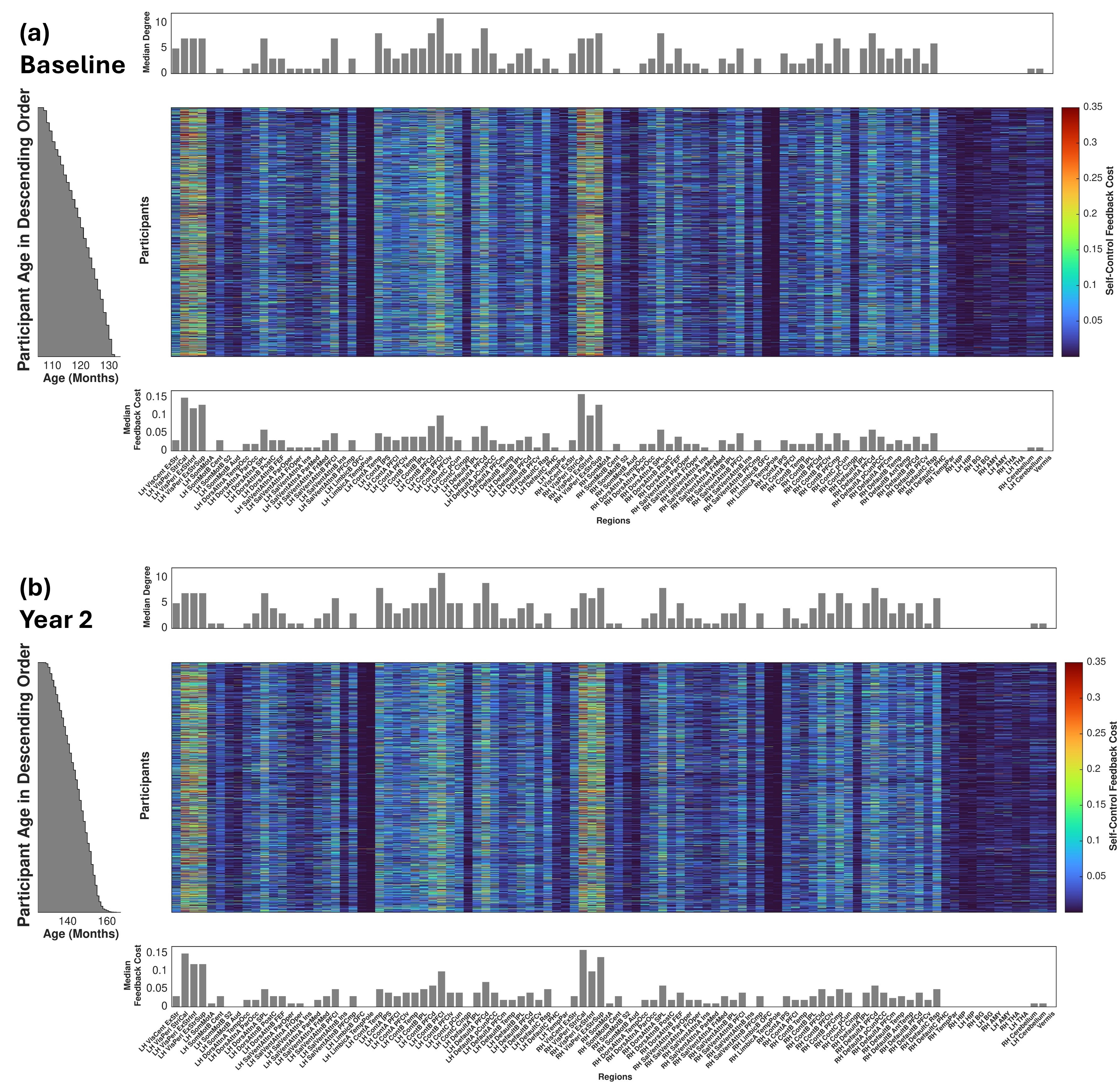

**Figure S1:** Feedback costs at which a region’s control action only depends on its own dynamic state, as a function of participants (y-axis) and analyzed regions (x-axis). The color range corresponds to cost values, estimated from the best-quality run at baseline (a) and two-year follow up (b). Participants are sorted by age. Median (over participants) regional connectedness (degree) for each analyzed region is shown in the bar plot above each heat map, and median feedback costs in the bar plot below each heat map. All parameters were estimated from adjacency matrices obtained by thresholding corresponding connectivity matrices based on the moderate outlying peak cross-correlation value.

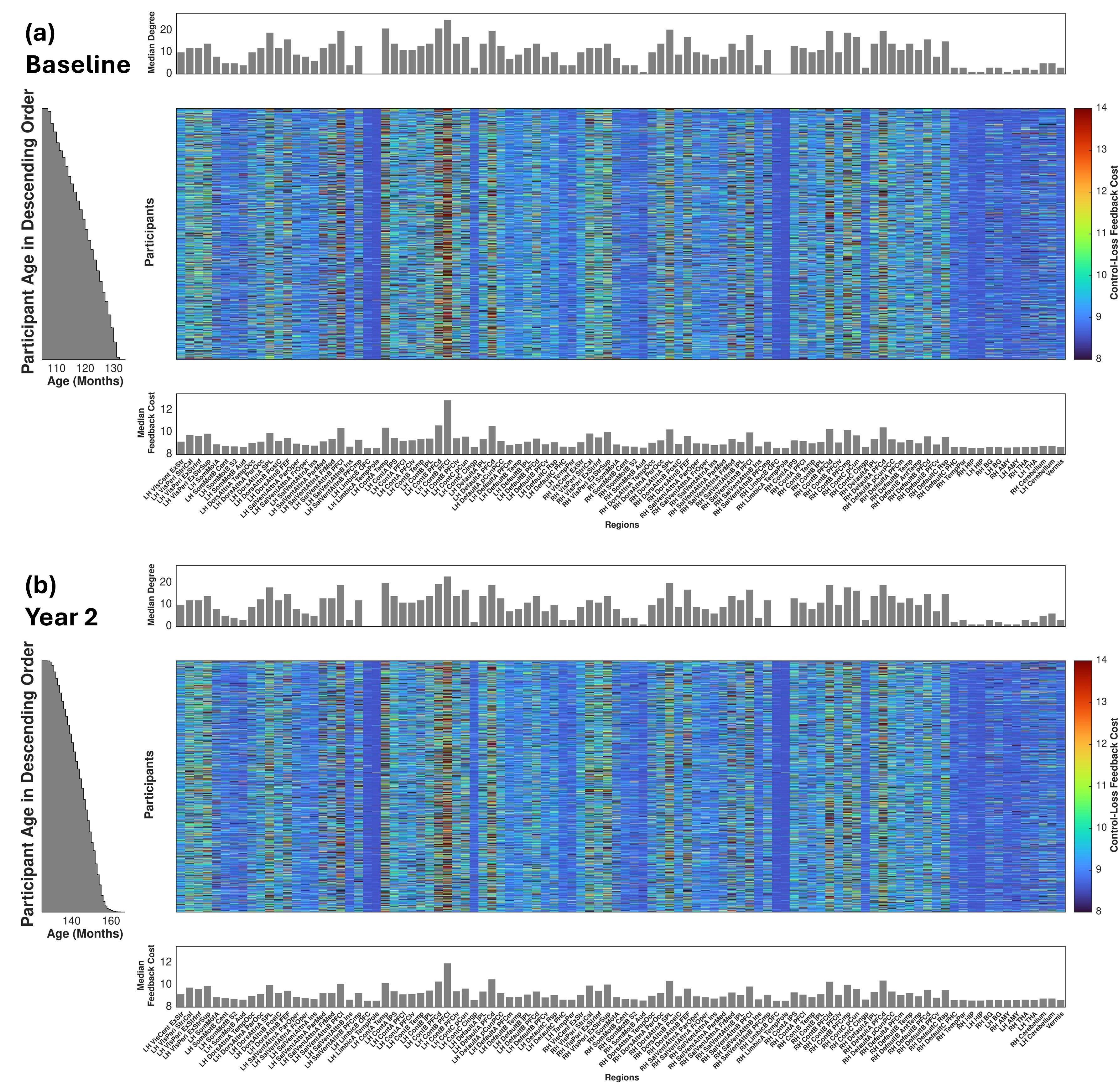
**Figure S2**: Feedback costs at which regions lose their control action on the network, as a function of participants (y-axis) and analyzed regions (x-axis). The color range corresponds to cost values, estimated from the best-quality run at baseline (a) and two-year follow up (b). Participants are sorted by age. Median (over participants) regional connectedness (degree) for each analyzed region is shown in the bar plot above each heat map, and median feedback costs in the bar plot below each heat map. All parameters were estimated from adjacency matrices obtained by thresholding corresponding connectivity matrices based on the 75^th^ percentile of peak cross-correlation value.

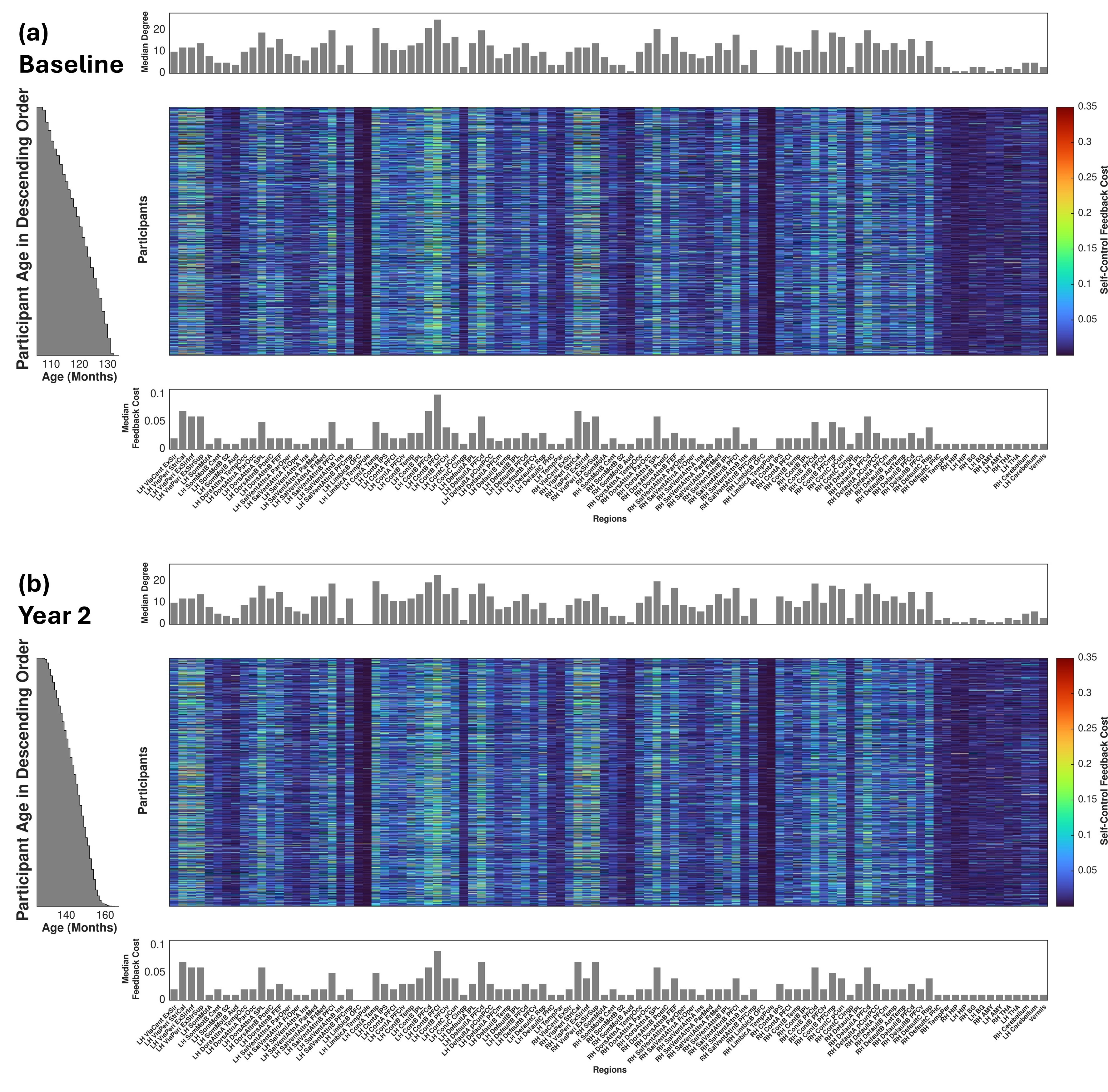

**Figure S3**: Feedback costs at which a region’s control action only depends on its own dynamic state, as a function of participants (y-axis) and analyzed regions (x-axis). The color range corresponds to cost values, estimated from the best-quality run at baseline (a) and two-year follow up (b). Participants are sorted by age. Median (over participants) regional connectedness (degree) for each analyzed region is shown in the bar plot above each heat map, and median feedback costs in the bar plot below each heat map. All parameters were estimated from adjacency matrices obtained by thresholding corresponding connectivity matrices based on the 75^th^ percentile of peak cross-correlation value.

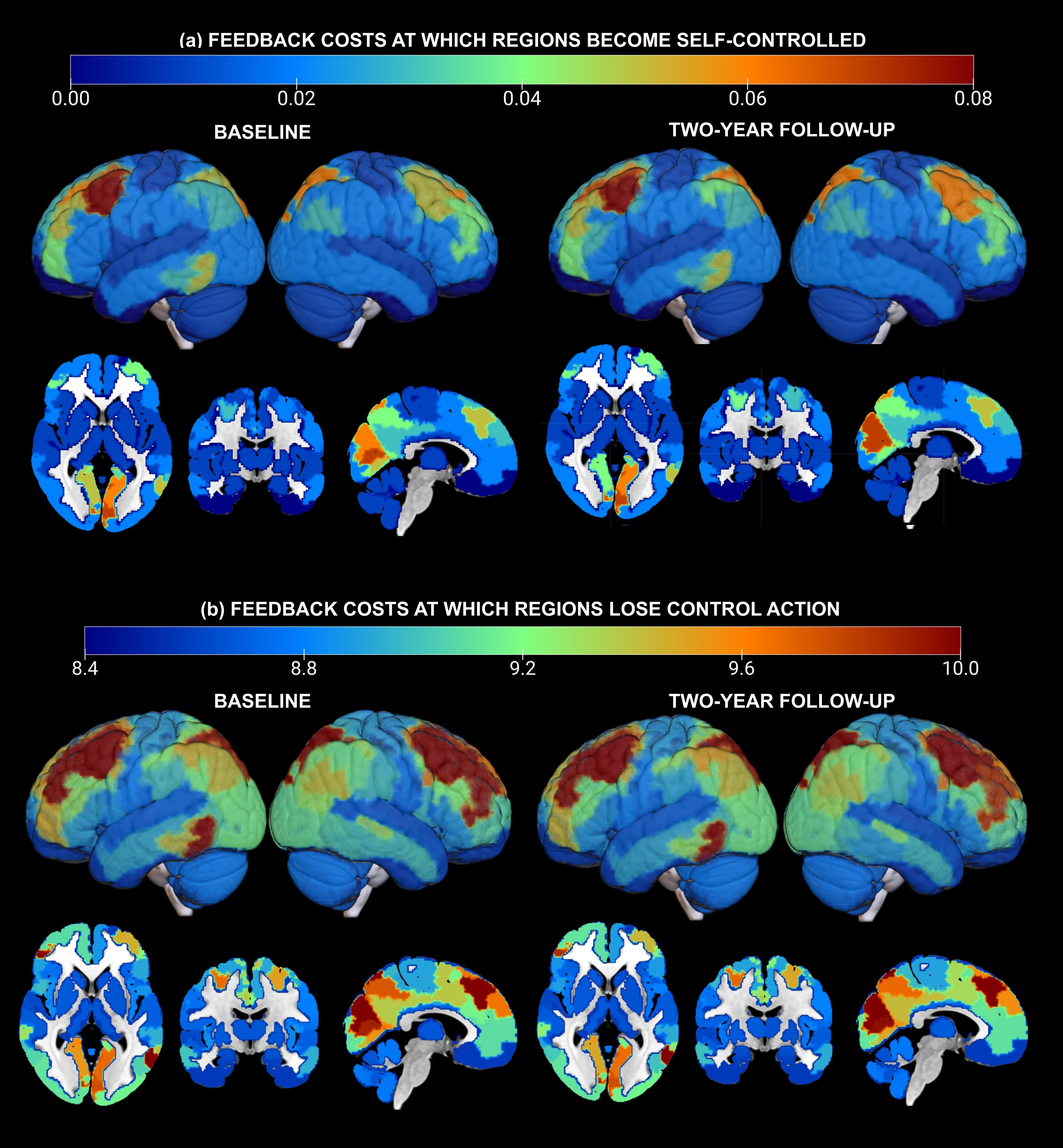
**Figure S4**: Spatial distribution of median (across participants) feedback costs at which (a) a region’s control action depends only on its own dynamic state, and (b) a region loses its control action over the network. Distributions are shown separately for baseline (left) and two-year follow up (right), and two-dimensional (horizontal, coronal, sagittal) slices and three-dimensional volumes are superimposed. Colors correspond to feedback cost values. All parameters were estimated from adjacency matrices obtained by thresholding the corresponding connectivity matrices based on the 75th peak cross-correlation value.

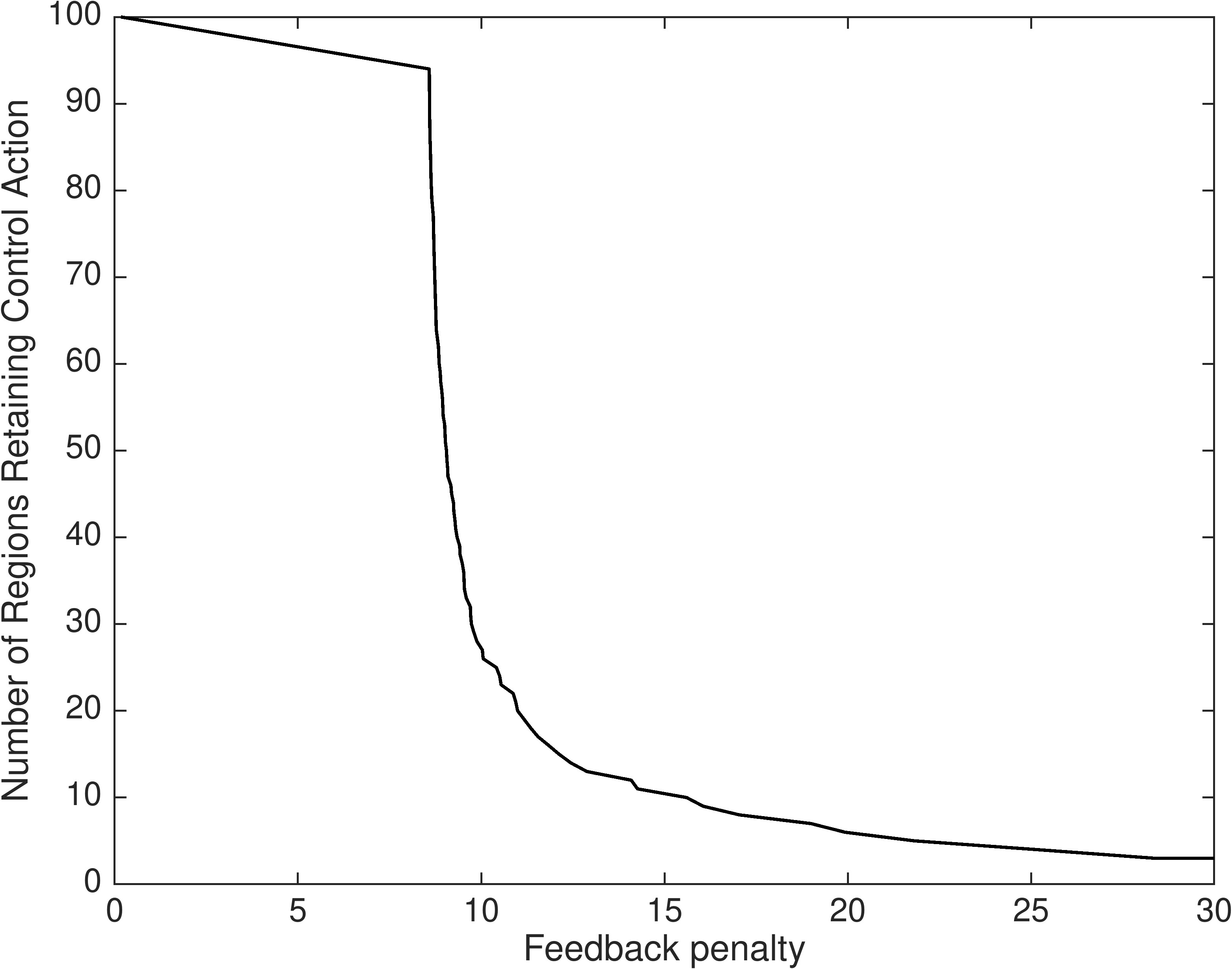

**Figure S5:** Example of number of regions that retain their control action as a function of increasing feedback cost (data are based on a representative brain from the analyzed sample).

| **FEEDBACK COSTS AT WHICH REGIONS BECOME SELF-CONTROLLED** | | |
| --- | --- | --- |
| **Demographic** | **Statistic** | **Value** |
| **Race-Ethnicity (0 = White Non-Hispanic,**  **1 = Others)** | Regions of negative correlations (-𝜷) | Bilateral visual striate calcarine  Left frontal eye fields  Left ventral lateral PFC  Left inferior parietal lobule  Right dorsolateral PFC |
|  | 𝜷* | [-0.0170, -0.0062] |
|  | 95% CI | [-0.0264, -0.0013] |
|  | P-Value | < 0.0365 |
| **Family Income** | Regions of negative correlations (-𝜷) | Left insula  Right primary somatomotor cortex |
|  | Standardized 𝜷* | [-0.0963, -0.0893] |
|  | 95% CI | [-0.1520, -0.0348] |
|  | P-Value | <0.0106 |
| **FEEDBACK COSTS AT WHICH REGIONS LOSE CONTROL ACTION** | | |
| **Demographic** | **Statistic** | **Value** |
| **Race-Ethnicity (0 = White Non-Hispanic,**  **1 = Others)** | Regions of negative correlations (-𝜷) | Left cuneus |
|  | 𝜷* | -0.1693 |
|  | 95% CI | [-0.2990, -0.0395] |
|  | P-Value | < 0.0212 |
| **Family Income** | Regions of positive correlations (+𝜷) | Right inferior parietal lobule |
|  | Standardized 𝜷* | 0.1037 |
|  | 95% CI | [0.0464, 0.1610] |
|  | P-Value | < 0.0035 |
|  | Regions of negative correlations (-𝜷) | Left medial PFC (part of the DMN) |
|  | Standardized 𝜷* | -0.0990 |
|  | 95% CI | [-0.1558, -0.0423] |
|  | P-Value | < 0.0063 |

**Table S1:** Statistics of mixed effects models testing associations between control costs (estimated from the best-quality fMRI run) and demographic characteristics. Reported p-values have been corrected for the False Discovery Rate (FDR).
